# High-affinity nanobodies as tools for structural and functional studies on mammalian Arc

**DOI:** 10.1101/2021.12.16.472929

**Authors:** Sigurbjörn Markússon, Erik I. Hallin, Helene J. Bustad, Arne Raasakka, Ju Xu, Gopinath Muruganandam, Remy Loris, Aurora Martinez, Clive R. Bramham, Petri Kursula

## Abstract

Activity-regulated cytoskeleton-associated protein (Arc) is a multidomain protein of retroviral origin with a vital role in the regulation of synaptic plasticity and memory formation in mammals. However, the mechanistic and structural basis of Arc function is little understood. Arc has an NTD involved in membrane binding and a CTD which binds postsynaptic protein ligands. In addition, the NTD and CTD both function in Arc oligomerization, including assembly of retrovirus-like capsid involved in intercellular signaling. We produced and characterised six ultra-high-affinity anti-Arc nanobodies (Nb). The CTD of both rat and human Arc could be crystallised in ternary complexes with two Nbs simultaneously bound (H11 and C11). H11 binding deep into the stargazing-binding pocket of Arc CTD suggested competitive binding with Arc ligand peptides, which was confirmed *in vitro*. This indicates that the H11 Nb could serve as a genetically-encoded tool for inhibition of endogenous Arc N-lobe interactions in study of neuronal function and plasticity. The crystallisation of the human Arc CTD in two different conformations, accompanied by SAXS data and molecular dynamics simulations, paints a dynamic picture of the mammalian Arc CTD. Dynamics were affected by mutations known to inhibit capsid formation, implying a role for Arc CTD dynamics in oligomerisation. Dimerisation of the NTD, together with structural dynamics of the CTD, suggest a mechanism, by which structural dynamics of the CTD may promote capsomer formation, and dimerisation of the NTD links capsomers, facilitating the formation of capsids. The described recombinant ultrahigh-affinity anti-Arc Nbs are versatile tools that can be further developed for studying mammalian Arc structure and function *in vitro* and *in vivo*.

## Introduction

Activity-regulated cytoskeleton-associated protein (Arc), is a highly conserved protein in vertebrates that has in recent years emerged as key regulator of long-term synaptic plasticity, with role in postnatal cortical development, memory, and cognitive flexibility. Arc is an immediate early gene in glutamatergic neurons that is highly expressed in a transient manner upon synaptic activation and salient behavioural experience [1]. Arc mRNA is directly targeted to dendrites following transcription, and translation occurs locally in activated synapses of glutamatergic neurons. The expression of Arc is dynamic, and the protein is quickly degraded [2–5]. Although loss of function studies have established causal roles for Arc, the molecular basis and cellular mechanisms of Arc are not fully understood [5, 6]. Arc serves as protein interaction hub, with several binding partners in the postsynaptic membrane and dendritic spines of excitatory synapses, as well as in the neuronal nucleus. Arc can also self-assemble, forming oligomers and large retroviral-like-capsids implicated in intercellular communication.

The induction of Arc expression is linked to the activation of NMDA receptors and activation of the mitogen-activated/extracellular-regulated protein kinase (MAPK/ERK) signalling cascade, both prerequisites of synaptic plasticity [7]. Furthermore, changes in the expression patterns of Arc are connected to changes in brain pathways linked to information processing [8], and the expression of Arc is upregulated during learning [9]. This suggests that Arc affects plasticity. Accordingly, dysregulation of Arc expression is implicated in plasticity-related diseases, such as fragile X syndrome and Alzheimer’s disease [10, 11], and Arc-deficient mice developed a severe deficiency in long-term memory formation [12].

Recent observations have further established the role of Arc as a fundamental regulator of synaptic plasticity. High expression levels of Arc led to the internalisation of AMPA receptors at the postsynaptic membrane [13, 14]. Furthermore, Arc directly interacts with both endophilin, dynamin, and clathrin-adaptor protein 2 (AP2) [15, 16]. Dynamin and endophilin both have essential late roles for clathrin-mediated synaptic receptor cycling, whereas AP2 has a critical role in initiating the process, as it coordinates cargo recruitment and selection by recruiting clathrin to the membrane [17]. A direct interaction of Arc with the AMPA receptor has been confirmed, as Arc binds the cytosolic tail of stargazin (Stg) [18]. Stg is an auxiliary subunit of AMPA receptors, associating with the receptor to modulate ion gating [19]. Arc has, therefore, been established as a primary regulator of AMPA receptors in the termination of LTP and LTD induction. Additionally, Arc interacts with inactive CaMKII, *i.e*. in the absence of calmodulin [20]. This interaction is thought to target Arc to weak synapses in LTP neurons, underlining its role in potentiation in long-term plasticity and reduction of AMPA receptors. This occurs in stimulated neurons in the terminal stage of LTP and in the initiation of LTD. As described above, LTP is often induced *via* the activation of NMDA receptors, and Arc co-localises with NMDA receptors in postsynaptic complexes [21]. While Arc requires postsynaptic density protein 95 (PSD95) for localisation to the postsynaptic density (described in more detail below), direct interactions with Glu2NA and Glu2NB have been identified [18, 22]. Glu2NA and Glu2NB are subunits of the NMDA receptor complex, vital for the induction of LTP [23]. Whether Arc regulates NMDA receptor function, or if it serves as an integrator between LTP and LTD, is not yet fully understood.

Mammalian Arc (mArc) consists of two folded domains, connected by a flexible linker, having disordered N- and C-termini. Structural and biochemical studies on the C-terminal domain (CTD) of Arc have shown it to consist of two separate four- and five-helix orthogonal bundles on the N- and C-terminal ends, respectively [18, 24]. The structure of the rat Arc CTD revealed a rigid bilobar structure, where the two lobes are connected by a short non-helical region [22]. The region following the CTD is fully disordered in solution, and it does not interact with the remainder of the domain [25]. The N-terminal lobe (N-lobe) of the domain harbours a buried peptide binding groove, which mediates many of the protein-protein interactions observed by Arc, including binding of Stg, GluN2a, GKAP and other peptides [18, 22, 24]. The binding pocket recognises a PxF/Y sequence motif, where the proline takes part in conserved C-H…*π* interactions, and the aromatic residue binds into a *π*-cluster within the hydrophobic core of the domain [24]. The Arc region preceding the peptide binding site, on the N-terminal end of the CTD N-lobe, is extended from the domain in the unbound state, but upon peptide binding, it folds against the bound peptide to form a two-stranded parallel *β*-sheet [22, 24]. In contrast, the C-terminal lobe (C-lobe) does not participate in peptide binding despite its structural similarity to the N-lobe.

In addition to the CTD, mArc contains an N-terminal domain (NTD) of uncertain function. The low solubility of the NTD has hindered structural characterisation. In contrast to the negative surface potential of the CTD, the NTD is highly basic. It was suggested to form a coiled coil, and using small-angle X-ray scattering (SAXS) and Förster resonance energy transfer (FRET), it was shown to pack against the CTD, resulting in a relatively compact fold of full-length Arc [26]. The NTD is required for the association of Arc with endophilin [15], although direct binding has not been demonstrated. Fulllength Arc, but not the CTD, associates with membranes enriched in anionic lipids, suggesting a role for the NTD in membrane association [26]. Membrane association of Arc can introduce negative curvature in anionic membranes [27], and the NTD contains a cysteine cluster (^94^CLCRC^98^), which is S-palmitoylated *in vivo*, likely to facilitate membrane binding [28].

Transposable elements (TEs), such as retrotransposons, constitute a major component of vertebrate genomes that can cause deletions and genomic instability. However, depending on the genomic context at their insertion site, they can sometimes be positively selected for in an exaptation, or domestication, event. Arc, likely introduced into the tetrapod genome *via* retroviral insertion, was initially identified in a search for Gag homology proteins descendant from Ty3/Gypsy retrotransposons [29]. Structural studies on the Arc-CTD show striking similarity with capsid domains of retroviruses [18, 26, 30]. Arc retains its ancestral ability to form viral-like capsids, as it forms large capsids that encapsulate RNA to transfer it between neurons [31, 32]. Arc preferably forms these assemblies upon binding of its own mRNA [33]. In this way, Arc encapsulates its own mRNA and transports it between neurons in a viral infection-like manner.

Recent high-resolution structures of the capsids formed by *Drosophila melanogaster* Arc isoforms 1 and 2 (dArc1 and dArc2, respectively) revealed assemblies of icosahedral symmetry with 240 Arc protomers [34]. There are, however, fundamental differences in the structure of mArc and dArc. dArc lacks the coiled-coil NTD found in mArc, and the two likely originated from two separate domestication events, suggesting evolutionary convergence [32]. Both dArc isoforms spontaneously assemble into capsids at ionic strengths mimicking RNA binding *in vitro* [34]. This is thought to occur in part due to the inherent tendency of the CTD lobes of dArc to oligomerise [35]. In contrast, the mammalian CTD is monomeric in solution [22, 24, 26], but dimeric forms of the CTD appear to be important in capsid assembly [36]. A larger role in the capsid assembly belongs to the NTD of mArc. Arc is unable to associate with mRNA and form higher-order oligomers in the absence of the NTD, and it has been suggested that full-length Arc is needed for the formation of the capsids, with the long linker region connecting the two domains facilitating domain swapping [26, 33]. Alanine scanning identified a motif within the NTD, ^113^MHVWREV^119^, as a main facilitator of capsid formation in rat Arc (rArc). Furthermore, a crystal structure of the motif revealed a dimeric coiled coil. Upon replacement of the motif by poly-Ala, the full-length protein loses higher-order oligomerisation and is dimeric [33]. The structure of the mArc capsid and the mechanism of capsid formation are unknown.

We produced and characterised six ultrahigh-affinity anti-Arc nanobodies (Nbs), single heavy chain antibodies derived from the variable heavy domain (VHH) of heavy chain camelid antibodies (HCAbs). Two of the Nbs led to crystal structures of the rat and human Arc-CTD, in an extended and collapsed conformation, visualising conformational dynamics of the Arc-CTD. The complementarity-determining region (CDR) 3 of NbArc-H11 bound deep into the peptide binding site of the Arc-CTD and inhibited peptide binding *in vitro*, suggesting applicability for studying Arc interactions at the molecular level. With complementary molecular dynamics (MD) simulations and SAXS data, we demonstrated the dynamic nature of the Arc-CTD. Moreover, we provide the first biophysical characterisation of the dimeric Arc-NTD. Taken together, the data suggest a novel mechanism of Arc capsid assembly, in which structural dynamics of the CTD facilitate capsomer formation, and NTD dimerisation leads to capsomer linking and formation of the mature capsid.

## Materials and methods

### Protein expression and purification

Fig. 1 outlines the Arc constructs. FLrArc-7A, full-length rArc containing a poly-Ala mutation (s113-119), was in the pHMGWA vector [33], resulting in an N-terminal His6-maltose binding protein (MBP) fusion [37]. *E. coli* BL21(DE3)-RIPL cells, transformed with the pHMGWA-rArc-FLrArc7A construct, were used to inoculate 10-50 mL LB medium containing 100 μg/mL ampicillin and 34 μg/mL chloramphenicol, followed by overnight culture at +37 °C and 170 rpm. The starter cultures were diluted 100-fold into 1-3 L of the medium and outgrowth carried out at +37 °C and 200 rpm. When the cell density (OD600) reached 0.8-1.0, expression was induced with 1.0 mM isopropyl β-D-1-thiogalactopyranoside (IPTG) and maintained for 4 h at +30 °C and 200 rpm. Cells were harvested via centrifugation at 6,000 g and +4 °C for 25 min, and the pellet was resuspended in 20 mM HEPES pH 7.5, 100 mM NaCl, 1 mM DTT, 0.1 mg/mL hen egg white lysozyme (20 mL per 1 L expression culture). Lysis was carried out via a single freeze-thaw cycle and sonication (seven cycles of 3 x 25 W for 10 s and 0 W for 30 s) and the soluble fraction harvested via centrifugation at 30,000 g and +4 °C for 45 min. The soluble fraction was extruded through 0.45-μm syringe filters and supplemented with 20 mM imidazole pH 7.5 and 1 mM dithiothreitol (DTT) before loading on a Ni^2+^-nitrilotriacetic acid (NiNTA) matrix equilibrated in 20 mM HEPES, 150 mM NaCl, 20 mM imidazole, 1 mM DTT, pH 7.5. The column was washed with 12 column volumes of the same buffer, before eluting the bound protein with the purification buffer containing 300 mM imidazole. To remove the His6-MBP tag, tobacco etch virus (TEV) protease was added and the protein dialysed against 1 L of 20 mM HEPES, 150 mM NaCl, 1 mM DTT in 6-8 kDa molecular weight cut-off (MWCO) dialysis tubing at +4 °C overnight. To remove TEV protease, the cleaved fusion tags and undigested fusion proteins, the protein was subjected to negative NiNTA purification following the same procedure as above but omitting imidazole from the washing buffer. The flowthrough and wash fractions were subjected to further negative purification using amylose resin in the same buffer. All affinity steps were performed at +4 °C. The protein was concentrated to 2 mL in a 30 kDa MWCO spin concentrator and applied onto a HiLoad 16/600 Superdex 200 pg size exclusion chromatography (SEC) column (GE healthcare, IL, USA) equilibrated in 20 mM Tris-HCl pH 7.4, 150 mM NaCl. Purity of the SEC fractions was analyzed via SDS-PAGE, before pooling and concentrating to 10-20 mg/mL. The concentrated protein was split into 50 μL aliquots, snap-frozen in liquid N2 and stored at −80 °C. Pure proteins were not subjected to more than one freeze-thaw cycle before use in functional assays or crystallisation.

**Figure 1:**
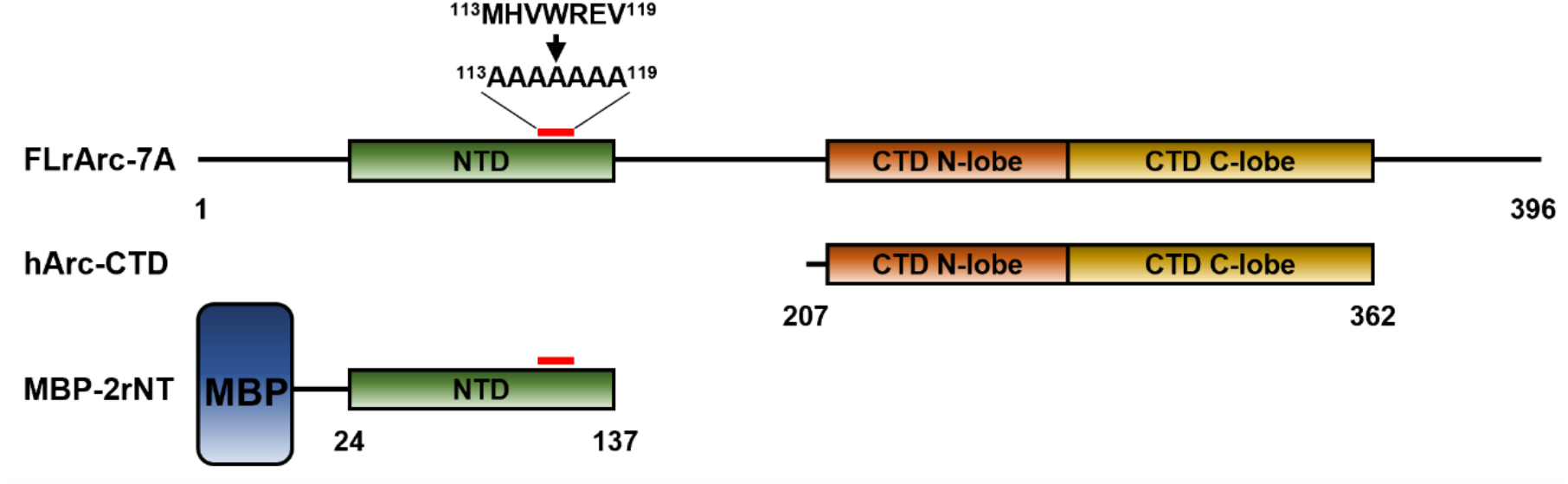
The Arc constructs used in this study. FLrArc-7A is the full-length rArc (residues 1-397) with the residues 113-119, in the second coil of the NTD mutated to hinder capsid aggregation [33]. hArc-CTD is the C-terminal domain (CTD) of the human Arc (residues 207-362). Not shown is the N-terminal 6xHis-tag and the 18-residue linker region containing the TEV site. 2rNT is the NTD of rat Arc (residues 24-137) containing the same poly-Ala mutation in its second coil. The domain proved to be insoluble in the absence of its fusion partner, maltose binding protein (MBP). FLhArc corresponds to FLrArc-7A, but with the human Arc sequence without any mutations.

The of human Arc CTD (hArc-CTD) (residues 206-361) was expressed and purified as described [26]. The protein was expressed with an N-terminal His6 tag in E. coli BL21(DE3) cells using the pTH27 vector and purified via affinity chromatography as described for the full-length protein above. As no cleavage of the His6 tag was observed upon incubation with TEV protease, the eluate from the first affinity step was concentrated to 2 mL in a 10 kDa MWCO spin concentrator and further purified on a HiLoad Superdex 75 16/600 pg SEC column equilibrated with 20 mM Tris-HCl pH 7.4, 150 mM NaCl. After SDS-PAGE analysis, the fractions of the main peak were pooled, concentrated to 15-30 mg/mL, aliquoted, snap-frozen in liquid N2 and stored at −80°C.

The NTD of FLrArc-7A (2rNT) was expressed as an N-terminal His6-MBP fusion construct in pHMGWA using *E. coli* BL21(DE3)-RIPL cells. A glycerol stock was used to inoculate 30 mL of LB medium containing 100 μg/mL ampicillin and 34 μg/mL chloramphenicol succeeded by overnight incubation at +37 °C and 170 rpm. The starter culture was diluted 100-fold into 3 L of the medium and incubated at +37 °C and 200 rpm. At OD600 = 0.5-0.7, expression was induced with 1.0 mM IPTG and maintained at +18 °C for 16-20 h. Cells were harvested via centrifugation at 6,000 g and 4 °C for 25-40 min, supernatant discarded and the pellet resuspended in 40 mM HEPES pH 7.5, 100 mM NaCl, 1 mM DTT, 0.1 mg/mL hen egg white lysozyme, supplemented with the cOmplete EDTA-free protease inhibitor cocktail (Roche, Basel, Switzerland) (25 mL of buffer per 1 L culture). Cells were lysed via a single freeze-thaw cycle and sonication and the soluble fraction collected via centrifugation at 30,000g and 4°C for 45 min. The soluble fraction was filtered through 0.45-μm syringe filters and applied to an NiNTA resin equilibrated in 40 mM HEPES, 400 mM NaCl, 20 mM imidazole, 1 mM DTT, pH 7.5 and the column washed in 15 column volumes of the same buffer. Bound protein was eluted with the same buffer containing 300 mM imidazole and directly applied to an amylose resin equilibrated in 20 mM HEPES, 400 mM NaCl, 1 mM DTT, pH 7.5, the column washed with 15 column volumes of the same buffer and eluted in 20 mM HEPES, 400 mM NaCl, 1 mM DTT, 10 mM maltose, pH 7.5. The protein was then concentrated to 2 mL in a 30 kDa MWCO spin concentrator and further purified on a HiLoad Superdex 200 pg 16/600 column equilibrated in 20 mM HEPES pH 7.4, 150 mM NaCl. Fractions assessed to contain pure fusion protein, via SDS-PAGE, were pooled and concentrated to 10-15 mg/mL. Glycerol to 10% (v/v) and 1 mM DTT were added to the concentrate before aliquoting, freezing in liquid N2 and storage at −80°C.

### Anti-Arc nanobody generation

Anti-Arc Nbs were obtained commercially from NanoTag Technologies (Göttingen, Germany). Briefly, two alpacas were immunized for a total of 6 times with the wild-type full length hArc and the FLrArc-7A mutant. Immunizations were started with wild-type hArc, and both proteins mixed in immunizations 3 to 5, gradually increasing the fraction of FLrArc-7A. Bleeding (150 mL) was carried out after immunization 6 and serum-based ELISA carried out to assess the total immune-response, using 50 ng FLrArc-7A and 1:100-1:32000 diluted serum. The initial Single domain antibody (sdAb) screening library was created by extracting total RNA from >150 mL blood from each animal and reverse transcribing to cDNA, using a 3-step nested PCR approach, thus specifically amplifying coding regions of IgG2 and IgG3 VHH fragments. The PCR product was cloned into a phagemid vector suitable for phage display and two rounds of phage display carried out, using FLrArc-7A as the target. SdAb sequences were extracted from the phages binding FLrArc, amplified and cloned into a screening vector, containing a C-terminal protease cleavable 3xFLAG-GFP tag. Lastly, 96 single colonies were analysed by ELISA using plates covered with 50 ng of FLrArc-7A per well. Detection was done with *via* the C-terminal 3xFLAG tag fused to the sdAbs. In addition, non-specific adsorption was assayed on plates coated with BSA as a negative control. Nearly all clones showed specific signals with an overwhelming majority of clones showing extremely high signals (>1.4), with only 4 clones testing negative. Following sequence analysis, sdAbs D4, C11, B5, H11, E5 and B12 were chosen as representatives of the four different sequence similarity families and two single binders and were obtained as His6-TEV-Nb constructs in prokaryotic pNT1433 expression vectors.

BL21(DE3) competent *E. coli* cells were transformed with the expression clones as outlined above. LB medium supplemented with 50 μg/mL kanamycin (10-20 mL) was inoculated with a single transformed colony, incubated overnight at 37°C and 170 rpm, diluted 100-fold into 0.5-2 L of the same medium and incubated at 37°C and 200 rpm. When OD600 reached 0.4-0.6, expression was induced via addition of 1 mM IPTG and maintained for 4 h at 30°C. Cells were harvested as described above and pellets resuspended in 40 mM HEPES pH 7.5, 100 mM NaCl, 20 mM imidazole, 0.1 mg/mL hen egg white lysozyme. Cells were lysed via a single freeze-thaw cycle and sonication and soluble fraction harvested via centrifugation at 30,000 g and 4°C for 30 min. NiNTA affinity purification and TEV proteolysis was carried out as above before a single negative NiNTA affinity purification step, concentration to 1-2 mL in 10 kDa MWCO spin concentrators and applied to either a HiLoad Superdex 75 pg 16/600 or a Superdex 75 Increase 10/300 GL SEC column (GE Healthcare, IL, USA) equilibrated in 20 mM Tris-HCl pH 7.4 and 150 mM NaCl. Fractions deemed pure via SDS-PAGE analysis were pooled, concentrated to 5-25 mg/mL, split into 50 μL aliquots, snap-frozen in liquid and stored at −80°C.

### Differential scanning fluorimetry (DSF)

To assess the thermal stabilisation of Arc constructs upon Nb binding, DSF was utilised. Proteins were diluted to 0.5-2 mg/mL in the assay buffer (20 mM Tris-HCl pH 7.4 and 150 mM NaCl) and mixed with 100x SYPRO-orange (in 50% (v/v) DMSO/assay buffer) in 384 well PCR plates to a final concentration of 5x, making the final DMSO concentration in the assay 2.5% (v/v). The final reaction volume was 18 μL. Fluorescence emission at 610 nm, following excitation at 465 nm, was measured in a LightCycler 480 LC RT-PCR system (Roche, Basel, Switzerland) over the temperature range 20-95 °C with a temperature ramp rate of 2.4 °C/min. Thermal denaturation midpoints (Tm) were determined as the maximum of the first derivative of the melting curves.

### Protein pulldown assays

For assessment of Nb binding and crude epitope mapping, protein pulldown assays were performed. 0.5 mg/mL of His-tagged hArc-CTD or His-MBP-tagged 2rNT were mixed with an equimolar amount of Nb and incubated on ice for 20-45 min. 200 μL of the solution were loaded on 100 μL of NiNTA agarose resin equilibrated in 20 mM Tris-HCl pH 7.4, 150 mM NaCl, 20 mM imidazole pH 7.5 and the mixture incubated at +4 °C under gentle agitation for 1 h followed by centrifugation at 200 g and +4 °C for 5 min. The supernatant (unbound protein) was removed, and the resin washed three times in the same buffer. To elute bound protein, the resin was incubated in the same buffer with 300 mM imidazole for 30 min before centrifugation as earlier. The contents of each fraction were analysed via SDS-PAGE.

### Isothermal titration calorimetry

The thermodynamics and affinity of Nb or peptide binding to FLrArc-7A were measured on a MicroCal iTC200 instrument (Malvern Panalytical, Malvern, UK). Binding of NbArcs to FLrArc-7A was measured in 20 mM Tris-HCl pH 7.4, 150 mM NaCl with 3.5-5 μM FLrArc-7A in the cell and 30-50 μM anti-Arc Nb in the syringe at 20 °C and a reference power of 10-12 μcal/s, with a stirring speed of 1000 rpm. In all cases, except for NbArc-E5, a single 0.5-μL priming injection was followed by 19× 2-μL 4 s injections of Nb with a spacing of 120 s and a filter period of 5 s. For NbArc-E5 binding to FLrArc-7A, the priming injection was followed by 38× 1-μL injections. In the case of stargazin (Stg) binding to FLrArc-7A, 2 mM of the Stg peptide (RIPSYRYR with N-terminal acetylation and C-terminal amidation) dissolved in the assay buffer was injected (1×0.5 and 19×2 μL) into 198 μM FLrArc-7A. For the Stg-Nb displacement assay, 150 μM of NbArc-H11 was injected (1×0.5 and 19×2 μL) into 10 μM FLrArc-7A and 120 μM Stg. Data processing (peak integration and dilution heat subtraction) was carried out in the Origin Lab software (v. 2021). Binding enthalpy (ΔH), association constant (Ka) and binding entropy (ΔS) were obtained by fitting the integrated thermograms with a 1:1 binding model.

### Small angle X-ray scattering

SAXS data from MBP-2rNT, and FLrArc-7A in the presence and absence of nanobodies, were collected on the SWING beamline of Soleil synchrotron, Saint Aubin, France in HPLC mode [38] using a wavelength (λ) of 1.03 Å and a sample-to-detector (EIGER 4X M) distance of 1.8 m, resulting in a momentum transfer (q) range of 0.005-0.5 Å^−1^ (q = 4π sin(θλ^−1^); 2θ is the scattering angle). In the SEC-SAXS setup, the samples run through a SEC column and into the capillary where X-ray scattering is measured. This method allows the collection of hundreds of scattering curves from the SEC peak that corresponds to the pure FLrArc-7A+Nb complexes and thus allows the removal of aggregates and unbound Nb. Nbs were mixed with FLrArc-7A in 1.4-fold molar excess, for a final complex concentration of 7 mg/mL prior to injection, and 50 μl of sample were injected onto a Bio SEC-3 300 column (Agilent Technologies) pre-equilibrated with the running buffer (20 mM HEPES, 150 mM NaCl, 0.5 mM TCEP, pH 7.5). The flow rate was 0.3 ml/min and data were collected with an exposure time of 990 ms and a dead time of 10 ms. Scattering of MBP-2rNT was measured in the same manner at a concentration of 8.3 mg/mL. Buffer data were collected at the beginning of the chromatogram and sample data were acquired in the peak area. Data reduction, R_g_ evaluation over elution profiles, data averaging and merging were performed using the beamline software Foxtrot (version 3.5.2).

SAXS data from hArc-CTD in the absence and presence of Nbs were collected on the BM29 beamline of European Synchrotron Radiation Facility, Grenoble, France in HPLC mode [39]. The data were recorded at a wavelength (λ) of 0.992 Å and a sample-to-detector (PILATUS 1M) distance of 2.867 m, covering a momentum transfer (q) range of 0.004-0.5 Å^−1^ (q = 4π sin θλ^−1^; 2θ is the scattering angle) [39]. Nbs were mixed with hArc-CTD in 1.3-fold molar excess for a final complex concentration of 10 mg/mL and 50 μl of the mixture injected onto an AdvanceBio SEC 130 column (Agilent Technologies) pre-equilibrated with the running buffer (20 mM HEPES, 150 mM NaCl, 0.5 mM TCEP, pH 7.5). Buffer data were collected at the beginning of the chromatogram and sample data were acquired in the peak area. Normalization and radial averaging were performed using the automated pipeline at the beamline [40].

Frame selection and buffer subtraction were carried out in CHROMIXS [41], primary analysis in PRIMUS [42] and distance distribution function (P(r)) analysis in GNOM [43]. *Ab initio* models were created using DAMMIN [44] and GASBOR [45]. Oligomer models were built from individual subunit models based on SAXS data in CORAL [46]. Theoretical scattering curves of coordinate files were calculated using CRYSOL [47] and scattering-based normal mode analysis of crystal structures carried out in SREFLEX [48].

### hArc capsid preparation and Nb association assay

FLhArc, full-length hArc, was cloned into the pETZZ_1a vector as a His-ZZ fusion protein construct with a TEV protease cleavage site [49]. *E. coli* BL21 CodonPlus cells transformed with the pETZZ_1a-FLhArc construct were used to inoculate 50 mL LB broth supplemented with 34 μg/mL chloramphenicol and 50 μg/mL kanamycin and incubated overnight at 28 °C and 200 rpm. The starter cultures were diluted into 1 L LB broth with antibiotics, and incubated at 37 °C and 200 rpm, until an OD_600_ of 0.6–0.8 was obtained. Expression was then induced by adding 0.5 mM IPTG and further incubated at 25 °C and 200 rpm overnight. Cells were harvested at 6000 × *g* for 30 minutes and pellets stored at −20 °C. Pellets were thawn on ice and resuspended in lysis buffer (5 mL/g pellet: 50 mM Na_2_HPO_4_/KH_2_PO_4_, 150 mM NaCl, 0.2 % Tergitol™ solution (Merck, Darmstadt, Germany), and 2 mM DTT, pH 8.0, containing 1 tablet cOmplete™ with EDTA (Roche, Basel, Switzerland), 10 mM benzamidine and 0.2 mM PMSF) and homogenised using a Thomas pestle tissue grinder, before sonication on ice for 3 × 45 s at 20 W, with 45-s pauses. The soluble fraction was harvested by centrifugation at 14 000 × *g* at 4 °C for 30 min. The soluble fraction was transferred to NiNTA agarose (Qiagen GmbH, Düsseldorf, Germany) equilibrated in 50 mM Na_2_HPO_4_/KH_2_PO_4_, 150 mM NaCl, and 2 mM DTT, pH 8.0, and incubated on rotation for 2 h at 4 °C. The matrix was then washed with 50 mM Na_2_HPO_4_/KH_2_PO_4_, 150 mM NaCl, 0.2% Tergitol™ (Merck, Darmstadt, Germany), and 2 mM DTT, pH 8.0, until A_280_=0, before washing overnight with at least 30 bed volumes of 50 mM Na_2_HPO_4_/KH_2_PO_4_, 150 mM NaCl, and 2 mM DTT, pH 8.0. The matrix was then washed 10 bed volumes of 50 mM Na_2_HPO_4_/KH_2_PO_4_, 1 M NaCl, and 2 mM DTT, pH 8.0 followed by 10 bed volumes 50 mM Na_2_HPO_4_/KH_2_PO_4_, 150 mM NaCl, and 2 mM DTT, pH 8.0, before elution with the same buffer containing 300 mM imidazole. The protein was dialysed against 50 mM Na_2_HPO_4_/KH_2_PO_4_, 150 mM NaCl, and 0.5 mM TCEP, pH 7.4 to remove imidazole, before cleaving off the fusion protein with TEV protease. To remove the HisZZ-tag, the cutting reaction was added to Talon (Takara Bio Inc., Kusatsu, Shiga, Japan) and gently agitated for 1 h at 4 °C, before elution and concentration at 2 000 × *g* at 4 °C. Concentration was determined using the theoretical extinction coefficient (Abs 0.1%) 1.71. A 260/280 ratio >1 indicated nucleic acid content, and capsid formation was checked by negative-staining electron microscopy.

For negative-staining transmission electron microscopy, 300-mesh copper grids (#Cu-300, Electron Microscopy Sciences, Hatfield, PA, USA) with formvar (#15820, Electron Microscopy Sciences) and carbon coating (provided by MIC, Department of Biomedicine, University of Bergen) were glow discharged for 1 minute at 30 mA in a Dieno pico 100 vacuum chamber (Diener Electronics). 5 μL FLhArc from the Nb association assay were applied to grids and incubated for 2 min before excess sample was removed using Whatman™ filter paper. Grids were then washed two times in 20 μL water droplets before they were stained with 2% uranyl acetate for 1 min, with excess solution removed with filter paper in between each wash and after staining. FLhArc without Nbs were used as control. Micrographs were obtained with a JEOL JEM-1230 electron microscope operated at 80kV. Images were recorded at 80,000× and 150,000× magnification.

For SEC analysis, 2 nmol of FLhArc were incubated with 1.5-fold molar excess Nbs and then applied to a Superdex 200 Increase 10/300 GL column (GE healthcare) equilibrated in PBS buffer.

### Protein crystallisation and structure determination

Protein crystallisation was carried out in TTP3 sitting drop 96-well crystallisation plates (SPT Labtech, Melbourn, UK) using a Mosquito LCP crystallisation robot (SPT Labtech, Melbourn, UK). For crystallisation of Nb complexes, Nb was mixed with the target protein in 1.2-1.5 fold molar excess and the complex isolated via SEC on a Superdex 75 Increase 10/300 GL or Superdex 200 Increase 10/300 GL (GE healthcare) column in 20 mM Tris-HCl pH 7.5, 150 mM NaCl. Fractions corresponding to the complex were pooled and concentrated to 10-37 mg/mL in 10 kDa MWCO spin concentrators and centrifuged at 20,000 g and 4 °C for 5-10 min before concentration determination *via* absorbance at 280 nm and set up of 96-well screening plates with 270-600 nL drops with varying protein:reservoir ratios (2:1, 1:1 and 1:2), sitting over a 70 μL reservoir of the precipitant solution. Plates were incubated at either 8 °C or 20 °C. 20 °C plates were automatically imaged in a Rock Imager-182 plate imaging system (Formulatrix, MA, US). Plates stored in an 8°C incubator were inspected manually under a microscope daily for the first week and weekly thereafter.

For matrix micro-seeding experiments, crystal seeds were produced by crushing protein crystals with a Pasteur pipette, end of which had been shaped into a ball using a flame. 4 μL of the reservoir solution were pipetted onto the crystal fragments and moved into a Seed Bead Eppendorf tube (Hampton research). This was repeated until the accumulated volume reached 28 μL, to create the undiluted seed stock. The seed stock was vortexed in 10 s intervals for 1-2 min and serially diluted four-fold to a 4,096x dilution. Crystallisation *via* seeding was carried out in 48-well hanging drop plates, with 2 μL drops, consisting of 1 μL protein, 0.75 μL reservoir and 0.25 μL of the appropriately diluted seed stock, over 200 μL reservoirs.

Diffraction data were processed in XDS [50] and scaled in XSCALE [50]. Analysis of crystal defects was carried out in XTRIAGE [51] and phases solved using molecular replacement (MR) in PHASER [52]. Refinement was carried out in PHENIX.REFINE [51] and manual model building in Coot [53]. XTRIAGE, PHASER and PHENIX.REFINE were all used within the Phenix software suite, v. dev-3958 [51]. Anisotropy analysis and anisotropic scaling was carried out using the STARANISO and the UCLA-DOE lab - Diffraction anisotropy [54] web servers. Structure validation was performed using MolProbity [55]. Analysis of Nb-epitope interfaces, crystal symmetry-based oligomers and interacting residues was carried out using the PDBsum [56] and PISA [57] web servers. All presented figures of crystal structures were prepared in PyMOL (v. 2.4.0., Schrödinger, LLC). Crystal diffraction and refinement statistics are shown in Table 1.

**Table 1:** X-ray diffraction data collection and refinement statistics. Statistics for the highest-resolution shell are indicated in parentheses.

|  | NbArc-E5 | rArc+H11+C11 | hArc+H11+C11,<br>extended | hArc+H11+C11,<br>collapsed |
| --- | --- | --- | --- | --- |
| **Data collection** |  |  |  |  |
| Beamline | I03 | P14 | P11 | P11 |
| Wavelength (Å) | 0.976 | 0.976 | 1.033 | 1.033 |
| Resolution range (Å) | 41.98-1.42 (1.46-1.42) | 38.51-2.70 (2.76-2.70) | 42.99-2.77 (2.84-2.77) | 48.00-1.94 (1.99-1.94) |
| Space group | P4<sub>3</sub>2<sub>1</sub>2 | P2<sub>1</sub>2<sub>1</sub>2<sub>1</sub> | P2<sub>1</sub>2<sub>1</sub>2<sub>1</sub> | P2<sub>1</sub>2<sub>1</sub>2 |
| Unit cell parameters | a=b=45.9 Å<br>c=103.75 Å<br>α=β=γ=90° | a=40.88 Å<br>b=92.81 Å<br>c=114.65 Å<br>α=β=γ=90° | a=40.80 Å<br>b=61.64 Å<br>c=171.95 Å<br>α=β=γ=90° | a=66.26 Å<br>b=139.29 Å<br>c=43.29 Å<br>α=β=γ=90° |
| Observed reflections | 201,623 (14,593) | 93,407 (5,153) | 77,407 (5,153) | 314,324 (6,893) |
| Unique reflections | 21,801 (1,581) | 12,476 (885) | 11,667 (849) | 26,403 (1,025) |
| Redundancy | 9.2 (9.2) | 7.5 (7.8) | 6.6 (6.0) | 11.9 (6.7) |
| Completeness (%) | 99.9 (99.8) | 99.2 (98.2) | 99.6 (99.9) | 86.5 (46.2) |
| Mean I/σ(I) | 9.5 (0.3) | 6.7 (1.0) | 7.5 (0.6) | 13.1 (0.8) |
| CC<sub>1/2</sub> (%) | 99.9 (33.3) | 99.1 (47.8) | 99.5 (32.4) | 99.9 (35.1) |
| R<sub>meas</sub> (%) | 10.8 (568.7) | 29.3 (255.2) | 25.1 (340.7) | 13.0 (235.6) |
| Mosaicity (°) | 0.143 | 0.121 | 0.618 | 0.113 |
| Wilson B-factor (Å<sup>2</sup>) | 19.43 | 36.55 | 48.96 | 30.95 |
| **Refinement** |  |  |  |  |
| R<sub>work</sub> (%) | 19.63 | 26.11 | 28.68 | 21.56 |
| R<sub>free</sub> (%) | 23.64 | 29.42 | 31.56 | 24.92 |
| RMSD<sub>bonds/angles</sub> (Å/°) | 0.005/0.722 | 0.018/2.210 | 0.005/0.757 | 0.008/0.775 |
| Ramachandran<br>favoured/allowed/outliers<br>(%) | 99.14/0.86/0.00 | 93.67/4.48/1.85 | 87.43/10.70/1.87 | 94.38/4.50/1.12 |
| Rama Z-score | -0.70 | -3.17 | -4.21 | -1.37 |
| Average B-factor (Å<sup>2</sup>) | 34.67 | 53.65 | 73.44 | 43.09 |
| **PDB code** | XXX | XXX | XXX | XXX |

### Crystallisation and structure solution of individual proteins and complexes

NbArc-E5. Anti-Arc Nb E5 was crystallised at 9.6 mg/mL under 1.26 M (NH4)2SO4, 0.1 M LiSO4, 0.1 M Tris-HCl pH 7.5 in a 2:1 300-nL drop over a 70 μL reservoir. Rod-shaped crystals in the tetragonal P4_3_2_1_2 spacegroup, with approximate dimensions of 300×40×40 μm^3^, grew after around 30 days at 20°C. Crystals were cryoprotected in the reservoir solution supplemented with 25% (v/v) glycerol. Diffraction data were collected at the I03 beamline at the Diamond light source (Oxford, UK) on an Eiger2 XE 16M detector at an X-ray wavelength of 0.976 Å (12.7 keV) with a 20×20 μm^2^ focused beam at 90.21% transmission. 1700 images were collected with an oscillation of 0.1° (for a total of 170°) with 50 ms exposures. Phases were solved using MR with 5HVG [58] as the reference model, for a translation function Z-score (TFZ) of 26.1.

rArc-CTD in complex with NbArc-H11 and -C11. A complex of the FLrArc-7A with Nbs H11 and C11 was prepared as described above, concentrated to 16 mg/mL and 150 μM Stg added to the concentrate prior to crystallisation. Needle-shaped crystals (200-400 μm in length) in the orthorombic P2_1_2_1_2_1_ spacegroup grew in 10% PEG10000, 10% PEG8000 after 5 months at 20 °C, in 1:1, 300-nL drop over a 70 μL reservoir. The crystals were cryoprotected in the reservoir supplemented with 25% PEG400. Diffraction data were collected at the EMBL/DESY P14 beamline of the PETRAIII synchrotron (Hamburg, Germany) on an EIGER 16M detector with an X-ray wavelength of 0.976 Å (12.7 keV). 2000 images were collected with an oscillation of 0.1° (for a total of 200°) and 10 ms exposure with an 150×61 μm^2^ flat beam at 100% transmission. As the crystals turned out to only contain the C-terminal domain of rArc in complex with the Nbs, phases were solved by initially obtaining a partial MR solution by using a crystal structure of the N-lobe of rArc-CTD in complex with Stg (4X3H; [18]), the C-lobe of rArc-CTD (4X3X; [18]) and a C11 homolog (6H16; [59]) as the reference models, giving a TFZ-score of 12.4. The final phases were obtained *via* MR, using a H11 homolog (4FHB, [60]) as the reference model and the partial solution from the earlier run, giving a TFZ-score of 17.4. The MR solution was subjected to automatic model building and density modification in AutoBuild within Phenix [51], followed by manual model building and refinement.

hArc-CTD in complex with NbArc-H11 and -C11. A complex of hArc-CTD and Nbs H11 and C11 was prepared as described above. Crystals with approximate dimensions of 150×50×50 μm^3^ in the P212121 space group grew from 37 mg/mL of the complex after around three weeks under 20% PEG3350, 0.2 Na malonate, 0.2 M KBr at 8 °C in a 1:1, 300-nL drop seated over a 70 μL reservoir. Crystals from the drop were cryoprotected in the reservoir solution supplemented with 25% PEG400. Diffraction data were collected at the P11 beamline [61] of the PETRA-III synchrotron (Hamburg, Germany) on a Pilatus 6M detector at an X-ray wavelength of 1.033 Å (12.0 keV). 2000 frames were collected with 0.1° oscillation (for a total of 200°) with a 20×20 μm2 focused beam at 2% transmission and an exposure time of 40 ms. Phases were solved via MR, using the hArc N-lobe in complex with Stg, C-lobe of hArc (6TNO and 6TN7, respectively;[24]) and the H11 and C11 Nbs from the rArc structure as the reference models, giving a TFZ-score of 17.8. The MR solution was run through AutoBuild [51], before manual model building and refinement. Efforts to solve the phases using the rArc-CTD-Nb crystal structure were unsuccessful.

hArc-CTD in complex with NbArc-H11 and -C11, collapsed crystal form. The ternary complex of hArc-CTD with Nbs H11 and C11 was prepared as described above and concentrated to 10.0 mg/mL. Microseeds of crystals of the same complex were prepared from crystals grown under 17% PEG3350, in a 1:1 300 nL drop, as described above. Crystals were grown using the hanging drop vapour diffusion method in drops consisting of 1 μL protein, 0.25 μL 1,024-fold diluted seed stock and 0.75 μL reservoir (12% PEG3350), hanging over 500 μL of the same reservoir solution. Needle shaped crystals in the orthorhombic P2_1_2_1_2 spacegroup grew after a 3-month incubation at 20°C. Crystals were cryoprotected in the reservoir solution supplemented with 25% PEG400. Diffraction data were collected on the P11 beamline at DESY [61], with an X-ray wavelength of 1.033 Å on a Dectris Eiger 16M detector. 3600 frames were collected with an oscillation of 0.1°/frame (360° total) using a 50×50 μm^2^ focused beam with an exposure time of 10 ms and 35% transmission. Due to an underestimation of the resolution limit upon collection, the data were collected with the detector edge at 2.4 Å. However, upon processing the data it became clear that the data extended much further and utilising the corners of the detector the data could be processed to 1.94 Å, but to lower than ideal completeness. Phases were solved using MR. Inititally, only the structure of H11 from the earlier hArc complex was used as the reference model, giving a TFZ score of 18.9, followed by the addition of the crystal structure of the hArc N-lobe in complex with Stg (6TNP; [24]), with Stg deleted from the model, for a TFZ score of 28.6. Finally, the C11 Nb structure from the earlier complex was used as the reference model, using a partial solution from the earlier run, to give a TFZ score of 31.9. Phases could not be solved using the C-lobe of hArc as a reference model, so what remained of the C-lobe was built into the model manually.

### Molecular dynamics (MD) simulations

Atomistic MD simulations were carried out on the hArc-CTD in GROMACS [62]. The hArc CTD crystal structure was stripped of Nbs, protons and ions, and assigned the predicted protonation state at pH 7.0 using PROPKA in PDB2PQR [63]. Protonated hArc-CTD was placed in a cubic box with a 10 Å extension around the protein. Solvation was carried out with the TIP3P water model and 0.15 M NaCl added to mimic physiological ionic strength. The models were subjected to conjugate gradient energy minimisation with a steepest decent step every 50th step and a maximum of 5000 steps. Temperature (NVT) equilibration to 300 K and pressure (NPT) equilibration, via isotropic pressure coupling, were carried out using the Berendsen thermostat. MD simulations were carried out using the OPLS-AA/L force field [64] and leap-frog algorithm, while retaining constant temperature and pressure using a velocity-rescale thermostat and a Parinello-Rahman barostat, respectively.

## Results

### Anti-Arc nanobodies and their interaction with FLrArc-7A

To aid in structural studies on Arc, Nbs were raised against the dimeric poly-Ala rArc mutant (FLrArc-7A). The initial llama immunization resulted in 92 positive Nb clones, and Nbs D4, H11, B5, C11, B12 and E5 were chosen as representatives of the six different CDR sequence families identified in the initial screen. Alignment of all the Nb sequences is shown in Fig. 2A. The alignment highlights the variable CDR loops, and demonstrates the unique nature of NbArc-H11, with an extended third CDR region. DSF was used to estimate thermal stability of the Nbs (Fig. 2B). No specific sequence trend could be identified, but as the framework region (FR) is highly conserved, the varying thermal stability likely depended upon the sequences in the paratope loops, mainly CDR3. Surprisingly, NbArc-H11 showed the highest T_m_ despite containing the longest CDR3, suggesting some folding of the CDR3 or its interaction with the conserved FR.

**Figure 2:**
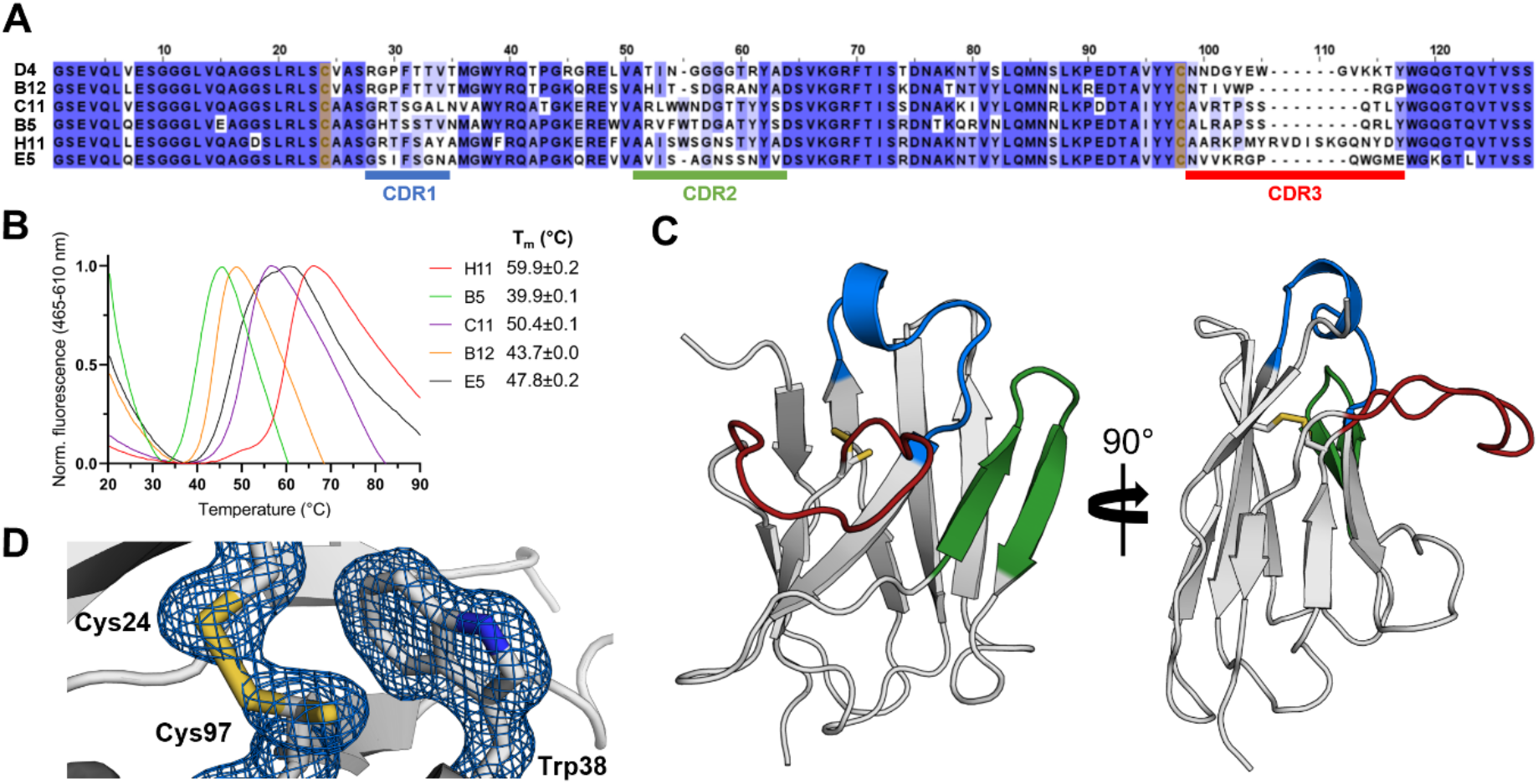
Anti-Arc nanobodies. **A** Sequence alignment of the six nanobodies reveals the location of the three CDRs, highlighting the long CDR3 of NbArc-H11. Sequences are coloured by conservation, and Cys24 and Cys92 (Kabat numbering) are highlighted in yellow. Sequence alignment was produced using the Clustal Omega web server [86] and visualised in JalView [87]. **B** Determination of NbArc T_m_ using DSF (N=3) in 20 mM Tris, 150 mM NaCl, pH 7.4 and 5x SYPRO™ orange, at a protein concentration of 1 mg/mL. The Tm of NbArc-D4 could not be determined. **C** 1.42 Å resolution crystal structure of NbArc-E5. Colouring of CDRs is the same as in panel *A*. **D** The conserved central disulphide bond, formed by oxidation Cys24 and Cys97, packs against Trp38 and appeared partially reduced. Electron density is shown as a blue mesh at 1.5σ.

To further characterise the Nbs in their free state, crystallisation was carried out. As NbArc-C11 and - E5 were the only ones initially obtained at sufficient yields and concentrations, crystallisation of other Nbs was not attempted. Crystallisation of C11 did not prove feasible. However, E5 crystallised and allowed for structure determination at 1.42 Å resolution (Fig. 2C). The structure shows the typical immunoglobulin fold, an incomplete *β*-barrel consisting of nine anti-parallel *β*-strands. The overall structure is compact, and the three CDR loops extend from the protein. Despite its elongated nature, the CDR3 loop is rigid in relation to the rest of the protein, as indicated by the refined B-factors (Fig S1A). Hydrophobic residues of the CDR3 pack onto the exterior of the central *β*-barrel to shield nonpolar side chains in a small hydrophobic core (Fig. S1B), which might account for the high solubility of this Nb. Furthermore, the two tryptophans and the basic side chains exposed in the CDR3 loop suggest a negatively charged, buried epitope. Surprisingly, the central disulfide of the protein (Cys24-Cys97), appeared mostly reduced, possibly due to radiation damage. Upon further inspection, it became clear that it was partially oxidised, and was refined to 20 and 80% occupancies for the oxidised and reduced rotamers of Cys97, respectively (Fig. 2D).

### Ultrahigh-affinity Nb binding to the CTD of Arc results in thermal stabilisation

To confirm the activity of the purified anti-Arc Nbs, and crudely map their epitopes, analytical SEC and protein pulldown experiments were performed (Fig. 3). Analytical SEC was chosen to quickly confirm binding to FLrArc-7A without the need for an affinity tag that might interfere with binding. The results (Fig. 3A) confirmed that all the purified Nbs bound Arc, as an increase was observed in the elution volume of the complexes when compared to the unbound protein alone. Although accurate molar mass and stoichiometry could not be determined from this experiment, the small shift in elution volume proposed the overall fold and the oligomeric state of the FLrArc-7A to remain unaltered upon Nb binding. To further determine whether the Nbs bound only one of the two domains of Arc, the His_6_-tagged hArc-CTD and His_6_-MBP fused 2rNT constructs were subjected NiNTA protein pulldowns with the Nbs (Fig. 3B). All the Nbs associated only with the CTD, but none bound to MBP-2rNT. This suggests that the NTD might have been inaccessible upon immunisation, possibly due to its position at the dimer interface of the protein.

**Figure 3:**
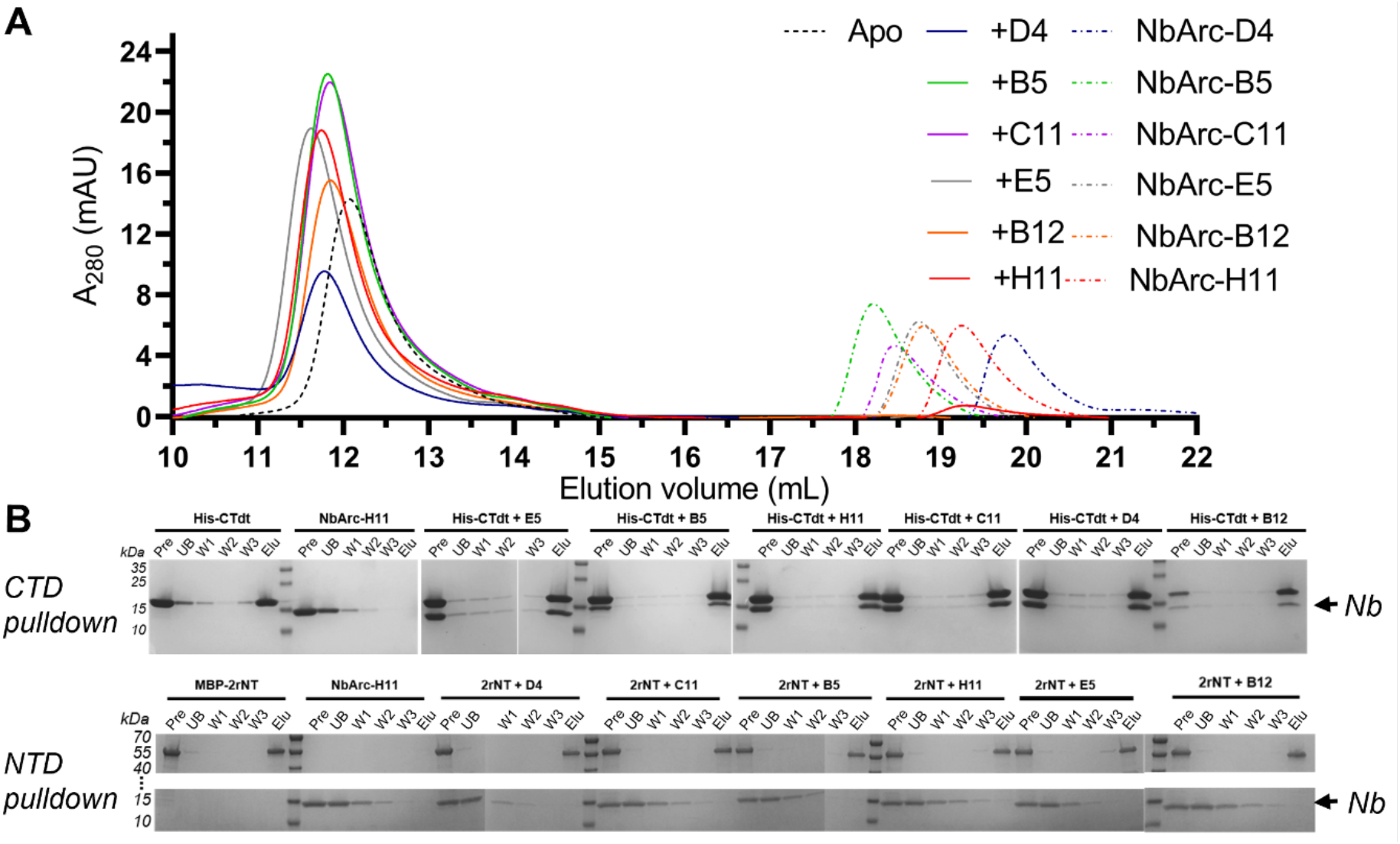
Initial characterisation of Nb binding to Arc. **A** Analytical SEC demonstrated Nb binding to FLrArc-7A. 2.2 nmol of FLrArc-7A were mixed with an equimolar amount of Nb, incubated on ice for around 30 minutes before the whole solution (100 *μ*L) was injected on an Superdex 200 Increase 10/300 GL SEC column in 20 mM Tris, 150 mM NaCl, pH 7.5. 2.2 nmol of the unbound protein (apo) and Nbs alone were run as controls. **B** NiNTA Nb pulldown assay with equimolar amounts of His-tagged hArc-CTD (CTD) or MBP-2rNT (NTD) and untagged Nbs, followed by SDS-PAGE analysis. All Nbs associated with the CTD (upper panel), as noted by the presence of the Nb band in all complex elutions, whereas none bound to the NTD (lower panel). Pre denotes the complex prior to loading on the matrix, UB the unbound fraction, W1-3 the three washing steps, and Elu the eluted fraction.

To study the thermal stabilisation of FLrArc-7A upon Nb binding, DSF was utilised (Fig. 4A, table 2). Binding of NbArc-C11 and -H11 resulted in highest stabilisation. This large increase in T_m_ suggested that these Nbs either stabilised flexible regions of Arc or selected for a conformer much more stable than unbound Arc. As all unfolding transitions observed were cooperative, and the measured T_m_ was in all cases higher than that of the individual components, it was assumed that the observed transition corresponded to the unfolding of the complex, except perhaps in the case of the rArc+B5 complex. Assessing the unfolding of the rArc+D4 complex proved unsuccessful, as two obscure transitions were observed, indicating complex dissociation prior to denaturation (data not shown).

**Figure 4:**
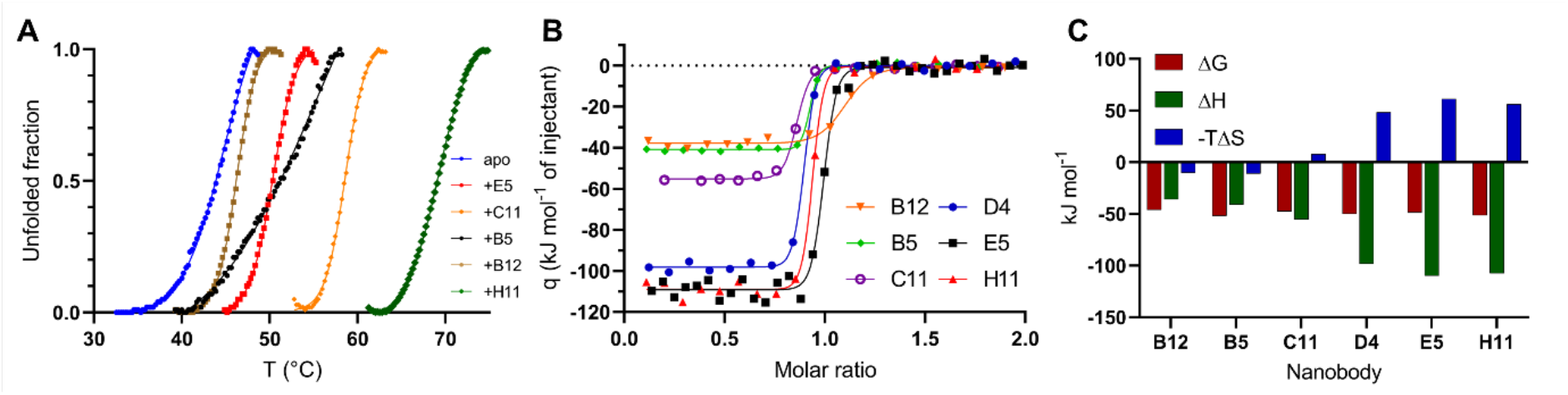
Characterisation of Nb binding to FLrArc-7A. **A** DSF melting curves of apo FLrArc-7A and in complex with Nb. Portions of the curves, other than the unfolding transition, are not shown for clarity. Tm values are shown in Table 2. **B** ITC titrations of Nb into FLrArc-7A. 30-50 *μ*M of Nb were titrated into 3.5-5 *μ*M of rArc at 20 °C in 20 mM Tris, 150 mM NaCl, pH 7.4. Raw ITC thermograms can be seen in Fig. S2. **C** Contributions of the enthalpy and entropy terms to the ΔG of Nb binding, highlighting the varying amount of enthalpy-entropy compensation.

**Table 2:** Thermal shift and thermodynamicsof Nb binding to FLrArc-7A. ΔT_m_ was measured using DSF (Fig. 4A). Thermodynamics of binding were derived from the integrated ITC isotherms shown in Fig. 4B. N/D: not determined.

| Nb | $\Delta T_m$ (°C) | Number of sites | $K_d$ (nM) | $\Delta H$ (kJ mol<sup>-1</sup>) | $-T\Delta S$ (kJ mol<sup>-1</sup>) | $\Delta G$ (kJ mol<sup>-1</sup>) |
| --- | --- | --- | --- | --- | --- | --- |
| B12 | +1.1 | 1.040±0.010 | 7.34±2.97 | -36.00±0.70 | -10.11 | -46.11 |
| B5 | +8.7 | 0.875±0.003 (~1) | 0.66±0.29 | -41.05±0.29 | -10.99 | -52.04 |
| C11 | +13.4 | 0.815±0.002 (~1) | 3.49±0.85 | -55.62±0.43 | 7.97 | -47.64 |
| D4 | N/D | 0.845±0.003 (~1) | 1.19±0.27 | -98.49±0.64 | 48.37 | -50.12 |
| E5 | +5.2 | 0.967±0.003 (~1) | 2.34±0.63 | -110.00±1.00 | 61.27 | -48.73 |
| H11 | +24.0 | 0.892±0.003 (~1) | 1.12±0.45 | -107.57±0.51 | 56.28 | -51.29 |

ITC was used to further assess the binding of the Nbs to FLrArc-7A (Fig. 4B-C, Table 2). All individual Nbs bound to Arc in 1:1 stoichiometry, in an enthalphy-driven manner. All bound with similar changes in Gibbs free energy (ΔG) of around −50 kJ mol^−1^, with estimated dissociation constants (*K_d_*) of 0.6 – 7.3 nM (Table 2). However, the affinity is possibly higher than that measured, as ITC is generally limited to affinities in the *K_d_* range of 10 nm – 100 μM [65], which was reflected in the large error margins of the *K_d_* estimations. Interestingly, the contributions of the enthalpy and entropy terms differed considerably (Fig. 4C). For NbArc-B12, -B5 and -C11, the entropy term was negligible, and the interaction was almost entirely enthalpy driven. However, NbArc-D4, -E5 and -H11 showed highly unfavourable binding entropy, compensated for by a large favourable enthalpy upon binding. For B12, B5 and C11, the negligible binding entropy suggested no considerable rigidification in the complex. However, the highly unfavourable binding entropy of D4, E5 and H11 suggested more rigidification, and increased bonding (hydrogen bonds and salt bridges) likely resulted in the compensating enthalpy. Altogether, ITC showed that the Nbs bind to FLrArc-7A with ultra-high affinity, and varying thermodynamics suggested varying modes of enthalpy-entropy compensation. Furthermore, the similarity of the binding thermodynamics of D4, E5 and H11, and B12, B5, ad C11, suggested that the six Nbs adopted only two binding modes upon complex formation.

### SAYS analysis of FLrArc-7A Nb complexes

SAXS was used to estimate the solution structure of the Nb complexes of FLrArc-7A, and to assess if any conformational changes occurred in Arc upon Nb binding (Fig. 5, Table 3). To measure scattering from only complexes and not unbound FLrArc-7A or Nbs, FLrArc-7A was mixed with excess Nb and subjected to SEC-SAXS, where the complexes were separated from the unbound FLrArc-7A using size-exclusion and SAXS frames collected as the proteins elute from the column. The scattering curves, and the derived parameters, indicate that in all cases the Nb bound to the FLrArc-7A dimer in 2:2 stoichiometry, confirming the Arc:Nb stoichiometry of 1:1 seen in ITC. The Kratky plot (Fig. 5C) indicated varying rigidity of the six complexes, without significant rigidification upon binding. The distance distributions (Fig. 5D) differed in all cases from that of unbound FLrArc-7A, showing an expansion in structure to a similar extent. R_g_ increased in all cases, accompanied by small changes in D_max_ (Table 3). The highest increase in R_g_, accompanied with an almost unchanged D_max_, was observed in complex with C11. This might indicate binding further from the centre of mass of FLrArc-7A. An increase in D_max_ was observed for E5 and H11 complexes, which indicated binding closer to the end of the longest axis.

**Figure 5:**
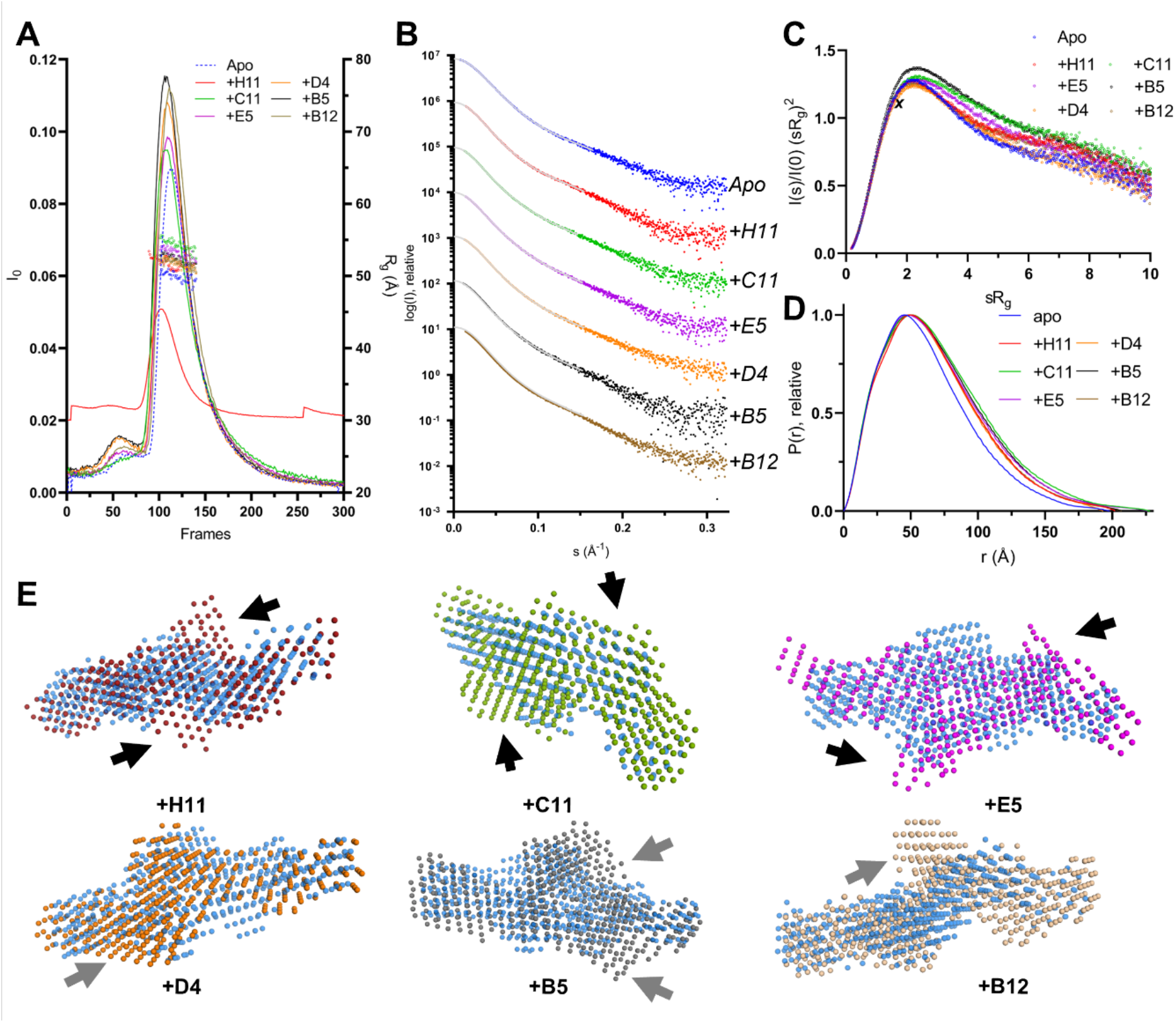
SEC-SAXS analysis of FLrArc-7A Nb complexes. **A** SEC-SAXS elution profiles of the complexes and unbound (Apo) FLrArc-7A. The estimated R_g_ of the frames used for processing is shown in open circles. All complexes were measured in 20 mM HEPES, 150 mM NaCl, 0.5 mM TCEP, pH 7.5. rArc+H11 was measured on a separate occasion in 20 mM Tris, 150 mM NaCl, pH 7.4, which caused differences in intensity and the elution volume of the main peak. **B** Scattering curves of rArc and rArc-Nb complexes, offset by one logarithmic unit for easier visualisation. Data fits from GNOM are shown as grey lines. **C** Dimensionless Kratky plot for all data sets. The maximum of an ideal rigid spherical particle (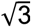, 1.104) is marked by ***X***. **D** Distance distribution profiles, obtained from the scattering data using GNOM. **E** *Ab initio* models of Nb-bound FLrArc-7A, shown at various orientations aligned with the model of the unbound protein (in transparent blue). Arrows indicate additional volumes observed in the complex models, which might correspond to bound Nb. Black arrows indicate two-fold symmetric binding, with the arrows rotated around the apparent symmetry axis, and grey arrows indicate additional volumes following no specific symmetry. Models were produced in DAMMIN with no forced symmetry, with *χ^2^* values of 0.992 (apo), 1.063 (+H11), 1.090 (+C11), 1.106 (+E5), 1.287 (+D4), 1.090 (+B5) and 1.090 (+B12).

**Table 3:** Parameters derived from SAXS of FLrArc-7A Nb complexes. I_0_ denotes the extrapolated forward scattering intensity, R_g_ radius of gyration, D_max_ the longest dimension of the particle, and Q_P_ Mw the molar mass estimate produced using the Porod invariant, Q_P_.

| Complex | Apo | +B5 | +B12 | +C11 | +D4 | +E5 | +H11 |
| --- | --- | --- | --- | --- | --- | --- | --- |
| $I_0$ | 0.083 | 0.110 | 0.110 | 0.094 | 0.110 | 0.097 | 0.092 |
| $R_g$ (Å) | 48.16 | 51.97 | 51.67 | 54.18 | 51.39 | 52.38 | 51.48 |
| Porod volume (nm<sup>3</sup>) | 256 | 301 | 305 | 316 | 293 | 303 | 324 |
| $P(r) R_g$ (Å) | 50.57 | 54.75 | 53.62 | 56.63 | 53.80 | 56.03 | 55.47 |
| $D_{max}$ (Å) | 198.0 | 198.9 | 193.5 | 206.8 | 203.5 | 227.5 | 229.0 |
| Quality est. GNOM | 0.74 | 0.79 | 0.81 | 0.78 | 0.79 | 0.76 | 0.77 |
| $Q_P$ Mw (kDa) | 135.4 | 152.9 | 156.5 | 164.4 | 167.1 | 160.2 | 156.1 |
| 1:1 complex mass (kDa) | 45.0 | 58.2 | 57.7 | 58.1 | 57.9 | 57.8 | 59.1 |
| Oligomeric state/stoichiometry | Dimer | 2:2 | 2:2 | 2:2 | 2:2 | 2:2 | 2:2 |

*Ab initio* dummy atom models derived from the data and their comparison with the unbound FLrArc-7A are shown in Fig. 5E. Changes were in all cases observed in the calculated molecular envelope, and with NbArc-H11, -C11 and -E5 bound, additional volumes indicated the presence Nbs bound to each monomer. The changes in the overall shape of the protein may be related to reduced fluctuations in the intrinsically disordered regions of FLrArc-7A upon Nb binding. The SAXS experiments demonstrated that all Nbs bound the FLrArc-7A dimer in 2:2 stoichiometry, but the low-resolution nature of SAXS and the low molar mass of the Nbs compared to dimeric Arc did not allow for accurate epitope mapping.

### Anti-Arc nanobodies associate with high-molecular-weight hArc without promoting disassembly

Considering the reported ability of some Nbs raised against viral proteins to associate with capsids and facilitate deactivation [66, 67], investigating whether the same applied for the anti-Arc Nbs was of interest. The association of the Nbs to hArc capsids was assessed using analytical SEC (Fig. 6). Although a minor peak of a similar elution volume as Nb-bound, dimeric FLrArc-7A, appeared upon addition of the Nbs, most co-eluted with the high-molecular-weight forms of hArc in the void volume (Fig. 6B). Furthermore, capsid aggregates, like those present in hArc without Nbs were apparent in negative staining TEM images of the hArc-H11 complex (Fig. 6C-D). The dimer peak, not present in the absence of Nbs, was most prominent in complex with H11, suggesting some degree of Nb-induced capsid dissociation. The results suggested that the Nbs could be associated with the capsids, or other high-MW forms, but did not induce conformational changes that would lead to capsid disassembly. However, this does not necessarily exclude that Nb binding could inhibit the formation of Arc capsids.

**Figure 6:**
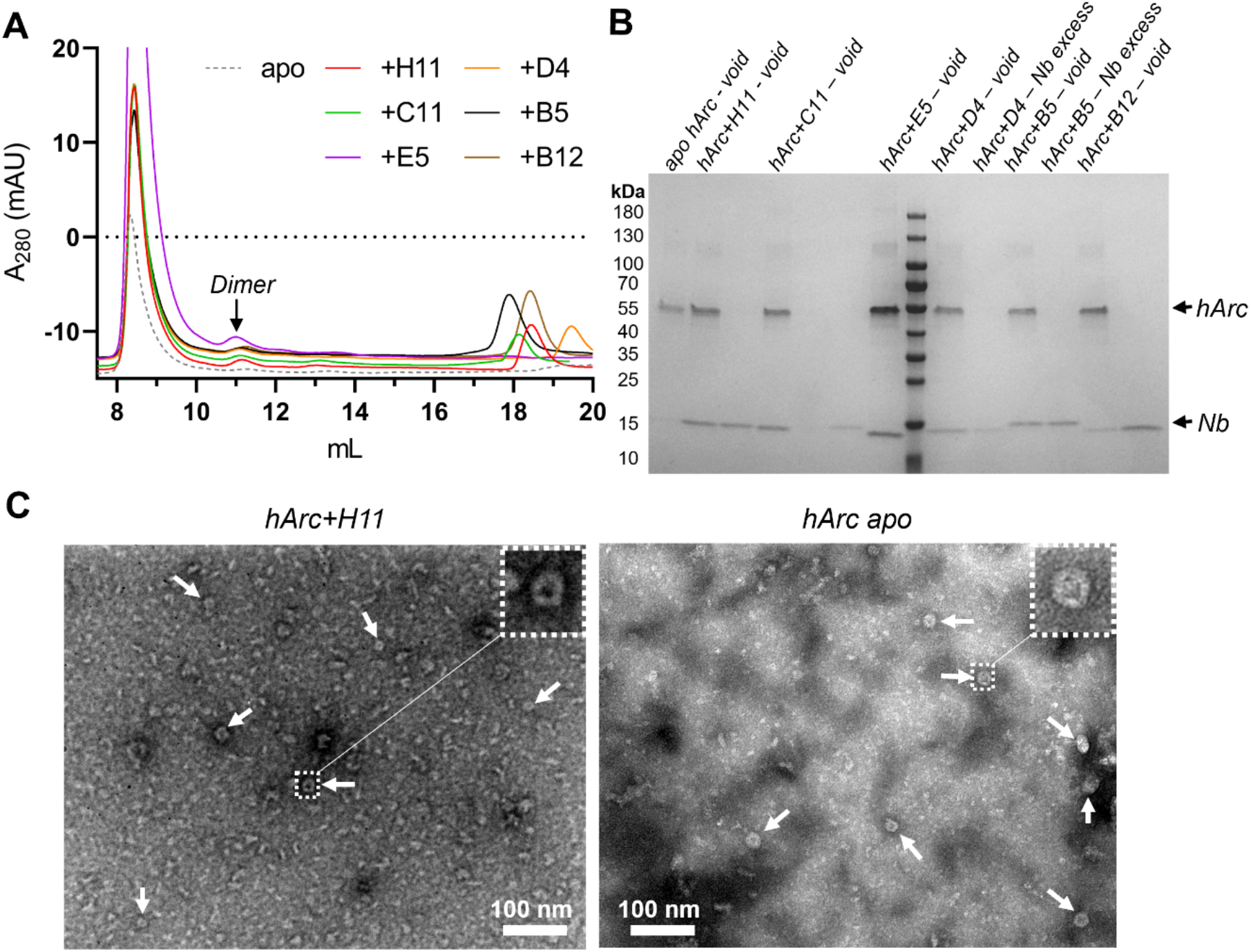
Rudimental analysis of Nb association with capsid hArc. **a)** Analytical SEC of capsid hArc-Nb complexes. 2 nmol of Arc capsids were mixed with a 1.5-fold molar excess of Nbs and run on a Superdex 200 Increase 10/300 GL in PBS buffer. Noted is the presumably dimeric peak present in some Nb complexes, with an elution volume corresponding to that of the FLrArc-7A complexes. NbArc-E5 was accidentally mixed with 4 nmol of hArc, resulting in the large void peak and lack of Nb excess peak. **b)** SDS-PAGE analysis of SEC fractions, demonstrating that Nbs elute with the capsids in the void peak. **c)** Negative staining TEM images of capsid hArc at 150,000x magnification in complex with NbArc-H11 (left) and without Nb (right). Arrows point at capsids, and a magnified views of selected capsids is shown.

### Crystal structure of an rArc-CTD ternary nanobody complex

A major aim of this study was the crystallisation of the FLrArc-7A mutant, using the six anti-Arc Nbs as crystallisation chaperones. However, despite extensive efforts, none of the complexes with one Nb grew crystals of sufficient quality for diffraction data collection. Therefore, crystallisation with more than one Nb bound was attempted. Analytical SEC showed the E5+C11, H11+C11, H11+B5 and E5+B5 Nb pairs to be compatible, whereas in other cases, binding of the second Nb caused dissociation of the first (data not shown). Crystallisation of FLrArc-7A was attempted in complex with both E5/C11 and H11/C11, as these individually led to the most thermal stabilisation upon binding (Fig. 4, Table 2). Whereas rArc+E5+C11 failed to crystallise, screens of FLrArc-7A in complex with H11 and C11 showed prominent crystal growth after extended incubation. The FLrArc-7A+H11+C11 crystals diffracted to 2.7-Å resolution, but the asymmetric unit was too small to contain full-length Arc in complex with both NbArc-H11 and -C11. Indeed, the crystals only contained the CTD of rArc in complex with both Nbs. This likely occurred due to proteolytic cleavage of the linker region.

Residues 211-356 of rArc, corresponding to the whole Arc-CTD, could be built into model, while the NTD was not detected in electron density. Nbs H11 and C11 could be fully built, only lacking the first two N-terminal and the last C-terminal residues (Fig. 7). The ternary complex shows the whole CTD of rArc, consisting of the N-lobe, a four-helix orthogonal bundle, and the C-lobe, a five-helix orthogonal bundle, connected by a straight rigid central *α*-helix (Fig. 7A). Nbs H11 and C11 bound to distinct, acidic epitopes on the N-lobe and the C-lobe, respectively. Accordingly, the electrostatic surface potential of the CDRs of both Nbs is basic (Fig. 7B). A large majority of crystal contacts were formed exclusively by the Nbs (Fig. S3A).

**Figure 7:**
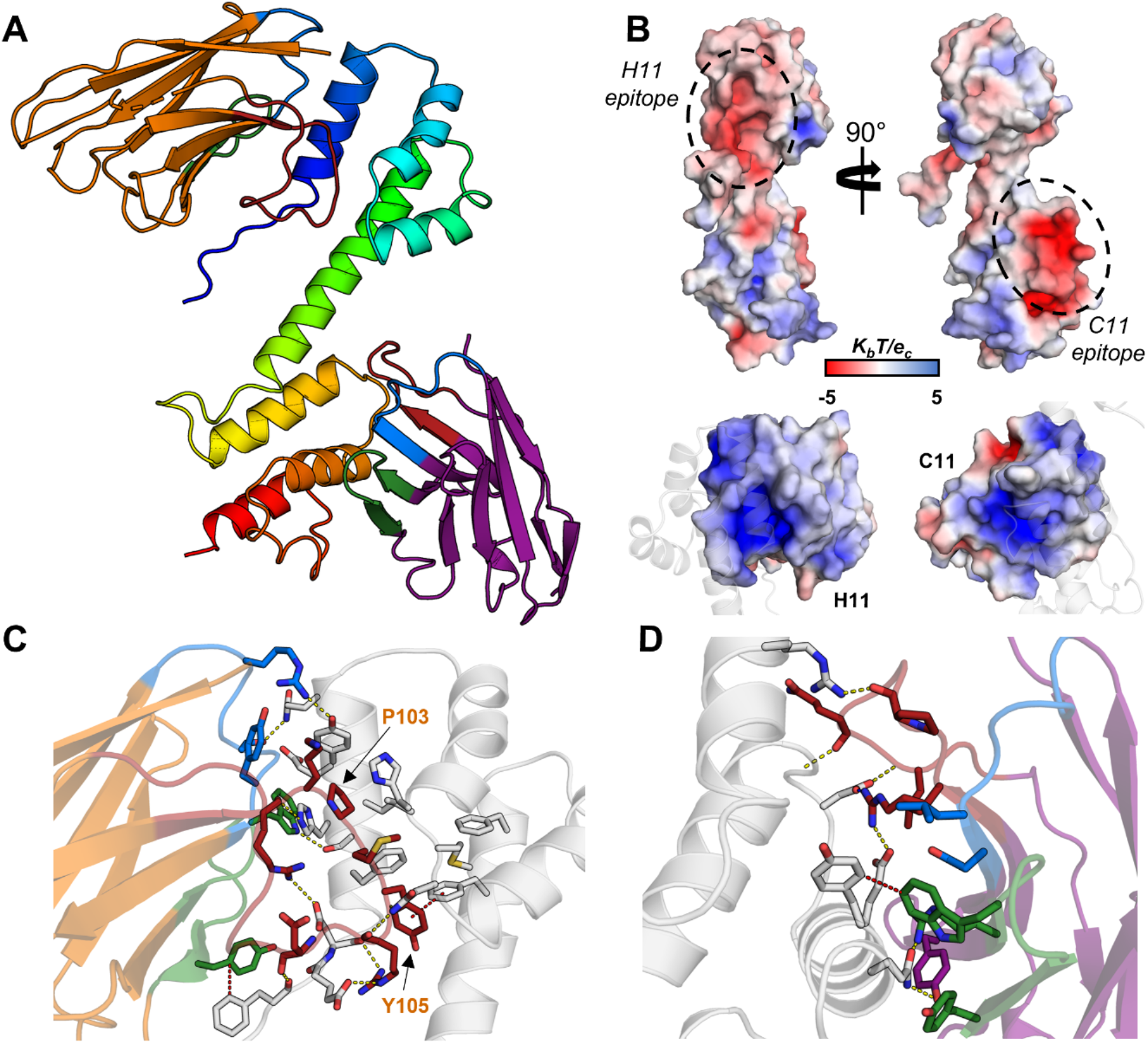
The 2.7 Å crystal structure of the rArc-CTD in complex with NbArc-H11 and -C11. **A** The overall structure of the ternary complex. The CTD of rArc is shown in blue to red from the N- to the C-termini. NbArc-H11 is shown in orange and NbArc-C11 in purple. CDR3 loops 1, 2, and 3 are colored blue, green and red, respectively. **B** Both Nbs bind acidic surface patches of the CTD (above), and the surface potential of the Nb CDRs is correspondingly positive (below). Electrostatic surface potential was calculated using ABPS [88]. **D** H11 binding to Arc Marked are Pro103 and Tyr105 of H11, which extend into the aromatic cluster of the N-lobe hydrophobic pocket. **D** C11 binding of the rArc-CTD. Polar contacts are indicated with yellow dashes and *π-π* interactions with red dashes. Colouring of the CDR loops in panels C and D is the same as in panel A.

NbArc-C11 bound a relatively solvent-exposed epitope, burying 738 Å^2^ of solvent-accessible Arc surface, and formed 2 salt bridges, 5 hydrogen bonds and 109 non-bonded contacts, including 3 *π-π* bonds (Fig. 7D). The size of the interacting area is as expected for antibody-antigen interactions, typically 600-800 Å^2^ [68]. In contrast, the elongated CDR3 loop of H11 protrudes from the β-barrel fold and extends into the N-lobe of the CTD. The binding buries a larger area on the CTD, 1044 Å^2^, and the first helix of the N-lobe is engulfed in a crevice between CDR loops 1 and 3 of H11 (Fig. 7C). H11 bound rArc through 2 salt bridges, 6 hydrogen bonds and 138 non-bonded contacts. CDR3 accounts for most of these interactions, and Pro103 and Tyr105 of CDR3 show CH…*π* and *π-π* interactions with aromatic residues within the hydrophobic core of the N-lobe. This groove of the N-lobe, coincidentally, is the ligand peptide binding site of Arc [18, 22, 24], indicating that nanobody H11 could be used to modulate protein-protein interactions of the Arc N-lobe. Additionally, H11 showed interactions with the N-terminal tail of the N-lobe, suggesting that it might affect the conformation of the long linker connecting the NTD and CTD in full-length Arc.

### Crystallisation of human Arc

Limited structural information has been available on hArc. The solution structure of the CTD of rat Arc had been determined [22], and despite the high sequence identity with hArc (only three conservative changes in the CTD; E318D, F334L and G359D), no structure of the intact hArc-CTD was available, apart from the individual N- and C-lobes [24]. Therefore, taking advantage of the opportunity presented by NbArc-H11 and -C11, crystallisation of hArc-CTD was carried out. The structure of the hArc-Nb ternary complex was refined to 2.77 Å resolution and contained the CTD of hArc (residues 209-355) in complex with the two Nbs (Fig. 8A). No density was observed for the N-terminal His6-tag. The structures of the two lobe domains are almost identical to those of the rat protein (Fig. 8B), with aligned Cα-RMSD of 0.65 Å for both the N- and C-lobes. However, the central helix connecting the two lobes breaks in hArc at residues 275-277, resulting in an altered relative orientation of the two lobes. Presumably, this was the cause of the altered unit cell, when compared to the rat Arc complex, with a shorter b-axis and an elongated c-axis (P2_1_2_1_2_1_; 40.88, 61.64, 171.95, 90, 90, 90), and the altered crystal packing where the bound Nbs had an even larger role in crystal contact formation (Fig. S3B).

**Figure 8:**
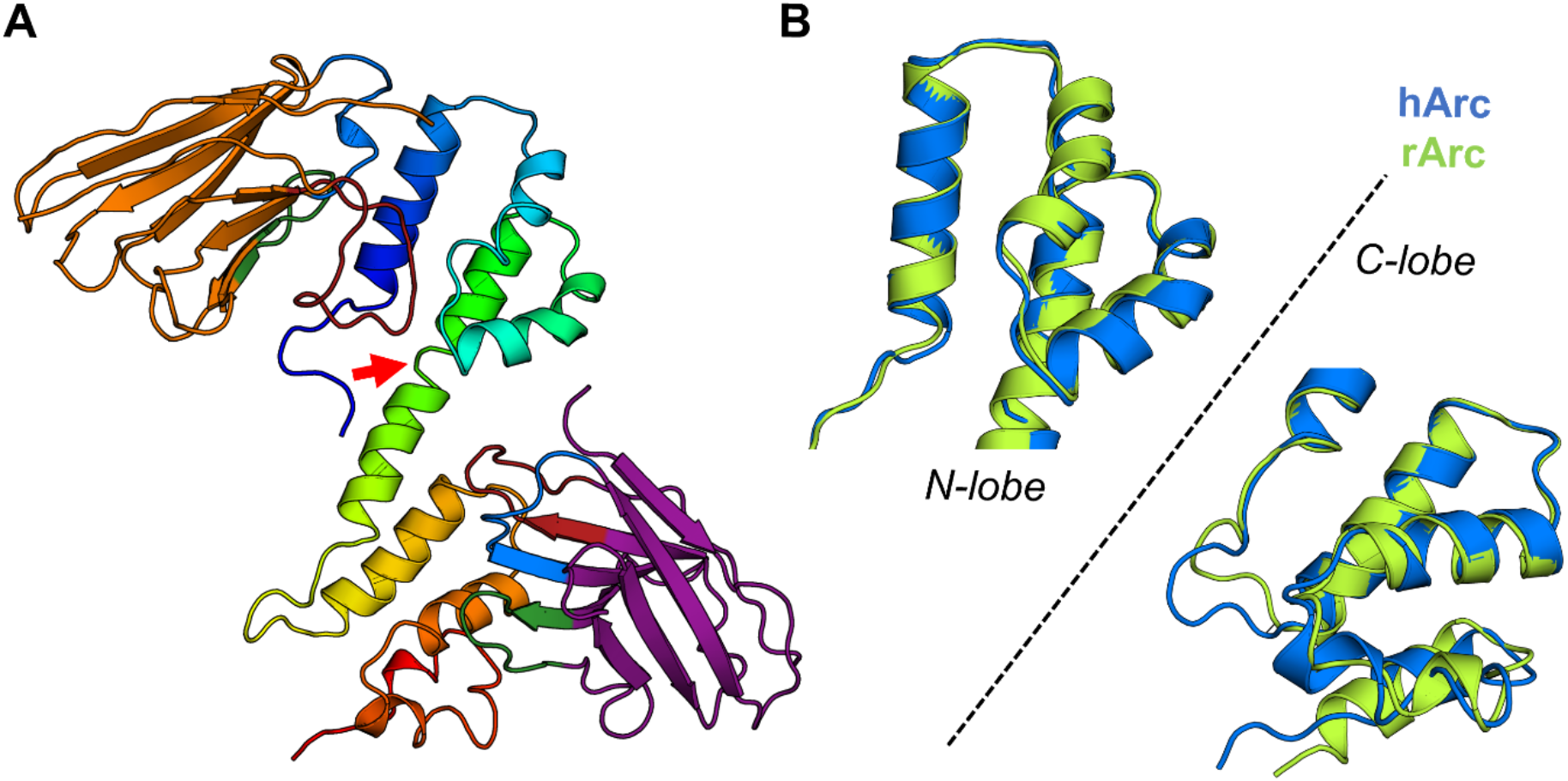
Crystal structure of the human Arc-CTD in complex with NbArc-H11 and -C11. **A** The structure of the ternary complex. The breaking of the central helix of the CTD connecting the two lobes is highlighted with a red arrow. The hArc-CTD is coloured blue to red from the N- to C-terminus, respectively, H11 is shown in orange and C11 in purple. CDR loops 1, 2, and 3 are coloured blue, green, and red, respectively. **B** Superimposition of the N-lobes and C-lobes of the rat (rArc) and human (hArc) Arc-CTD from the two crystal structures.

The binding mode of H11 to hArc was similar to that observed in the rArc structure, burying 1073 Å^2^ of solvent-exposed area on the CTD, with similar interactions. However, due to the bending of the central helix of the CTD, the binding by C11 was somewhat altered. The buried surface area of hArc was increased to 848 Å^2^, and binding was achieved *via* 3 salt bridges, 7 hydrogen bonds and 129 nonbonded contacts (including *π-π*, CH…*π* and van der Waals contacts). This was achieved by the additional interactions of C11 with both the central helix and the N-lobe, suggesting that the reduction of interactions in the rArc structure either occurred due to restrictions produced by crystal packing, or that C11 might select for a slightly bent conformer of the CTD by interacting with both lobes.

### NbArc-H11 inhibits ligand peptide binding of Arc

Arc localises to the postsynaptic density (PSD) and binds various ligand peptides into a hydrophobic groove located in N-lobe of the CTD. These peptides include Stg, GKAP, GluN2A, and WAVE1, involved in regulation of structural and functional synaptic plasticity [18, 22, 24]. This is the same site, to which the CDR3 loop of the H11 Nb bound in the crystal structures of both human and rat Arc. Upon further inspection, the conserved sequence motif of the ligand peptides is present in the H11-CDR3 loop, resulting in a nearly identical mode of binding (Fig. 9A and B). This suggested that H11, which binds with *K_d_* in the low nM range, could be used to efficiently displace bound peptides, which bind with affinities of 20-500 *μ*M [24], and inhibit peptide binding. To test this hypothesis, an ITC displacement assay was carried out, whereby the binding of H11 to Stg-bound FLArc-7A was measured (Fig. 9C, Table 4). FLrArc-7A bound the Stg peptide with a *K_d_* of 34.9 μM in an enthalpy-driven manner, corresponding well with previous obtained values for the isolated hArc N-lobe [24]. This demonstrated that the peptide binding affinity of the N-lobe was not altered in the full-length protein, confirming that the mutations in FLrArc-7A did not affect binding. Upon titration of H11 into Stg-bound FLrArc-7A, the measured binding enthalpy of around −75 kJ mol^−1^ corresponded to the binding enthalpy of H11 (−107 kJ mol^−1^) with the binding enthalpy of Stg (−39 kJ mol^−1^) subtracted. However, the ΔG of binding (and *K_d_*) of H11 binding to Stg-bound FLrArc-7A was unchanged, due to more favourable binding entropy. This indicated that what was measured was the binding of H11, accompanied by the dissociation of Stg, demonstrating that H11 facilitated displacement of the bound Stg peptide. Presumably, this also applies to other Arc ligand peptides which bind to the same site. Therefore, NbArc-H11 presents itself as a potential tool that could be used to inhibit various protein-protein interactions involving Arc *in vivo*, to study the function of these interactions in Arc-mediated regulation of synaptic plasticity.

**Figure 9:**
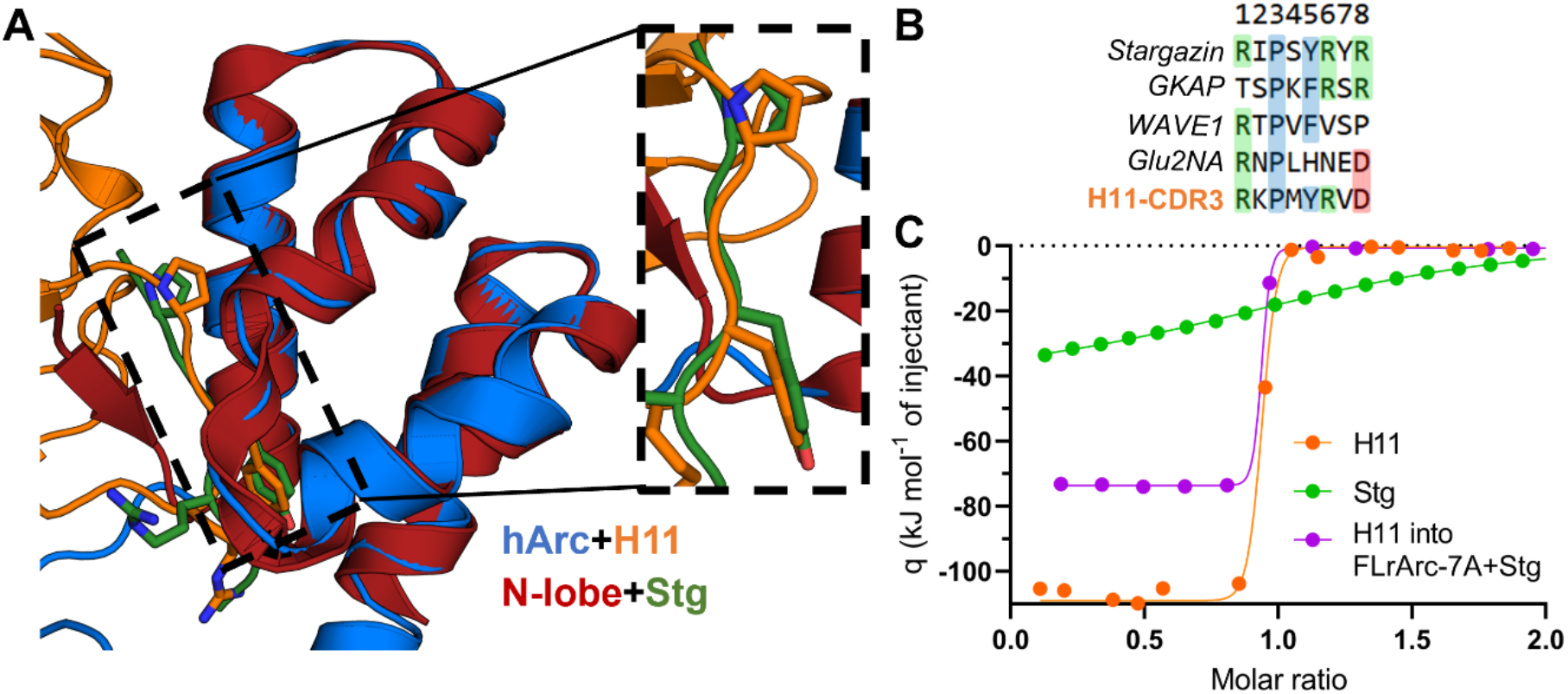
NbArc-H11 as a peptide binding inhibitor. **A** Alignment of hArc-CTD bound to H11 with the crystal structure of the human N-lobe with bound Stg (6TNP, [24]) reveals the identical mode of binding. **B** Sequences of various Arc ligand peptides aligned with the sequence of the H11-CDR3. **C** ITC displacement assay. Shown are the integrated thermograms of H11 and Stg titrated into FLrArc-7A, and H11 titrated into FLrArc-7A bound to Stg. The decreased binding enthalpy of H11 binding to Stg bound FLrArc-7A suggests that H11 binding leads to Stg dissociation. Raw thermograms are shown in Fig. S4. The H11 titration curve is the same as shown in Fig. 4B.

**Table 4:** ITC displacement assay to follow H11 and Stg binding to FLrArc-7A. The thermodynamic parameters of binding were obtained from the titration curves in Fig. 9C. Error margins indicate errors of data fitting from single experiments.

| Titration | Number of sites | $K_d$ | $\Delta H$ (kJ mol<sup>-1</sup>) | $-T\Delta S$ (kJ mol<sup>-1</sup>) | $\Delta G$ (kJ mol<sup>-1</sup>) |
| --- | --- | --- | --- | --- | --- |
| H11 | 0.892±0.003 | 1.12±0.25 nM | -107.57±0.51 | 56.28 | -51.29 |
| Stg | 1.090±0.011 | 34.90±0.2 μM | -38.50±0.64 | 15.54 | -22.96 |
| H11 into FLrArc-7A+Stg | 0.834±0.002 | 0.87±0.35 nM | -75.23±0.89 | 24.92 | -50.31 |

### Exploring the dynamic nature of the Arc-CTD central helix hinge region

The crystal structures of the human and rat Arc CTD showed almost identical folds, except in the conformation of the central helix. In the rArc structure, the helix is straight (Fig. 7), whereas in hArc, it is slightly kinked (Fig. 8), resulting in dislocation of the C-lobe in relation to the N-lobe. Upon comparison with the NMR model of the CTD from rArc [22], this conformational variation becomes even more apparent (Fig. 10A). Breaking of the helix at Gly277 results in a conformational shift, corresponding to a 3.7 Å translation of the C-terminal end of the helix in the hArc crystal and 14.4 Å in the NMR model (Fig. 10B). The dynamics of this helix might have a role in the function of Arc, and dynamic movements of the CTD could change between monomeric/dimeric Arc, with potential roles in synaptic plasticity, and the capsid form, which facilitates intercellular mRNA transport [5]. The corresponding helix in both dArc1 and dArc2 breaks, as the two lobes of CTD pack against each other in the capsid protomers [34]. To further explore this hypothesis, SAXS and MD simulations were carried out.

**Figure 10:**
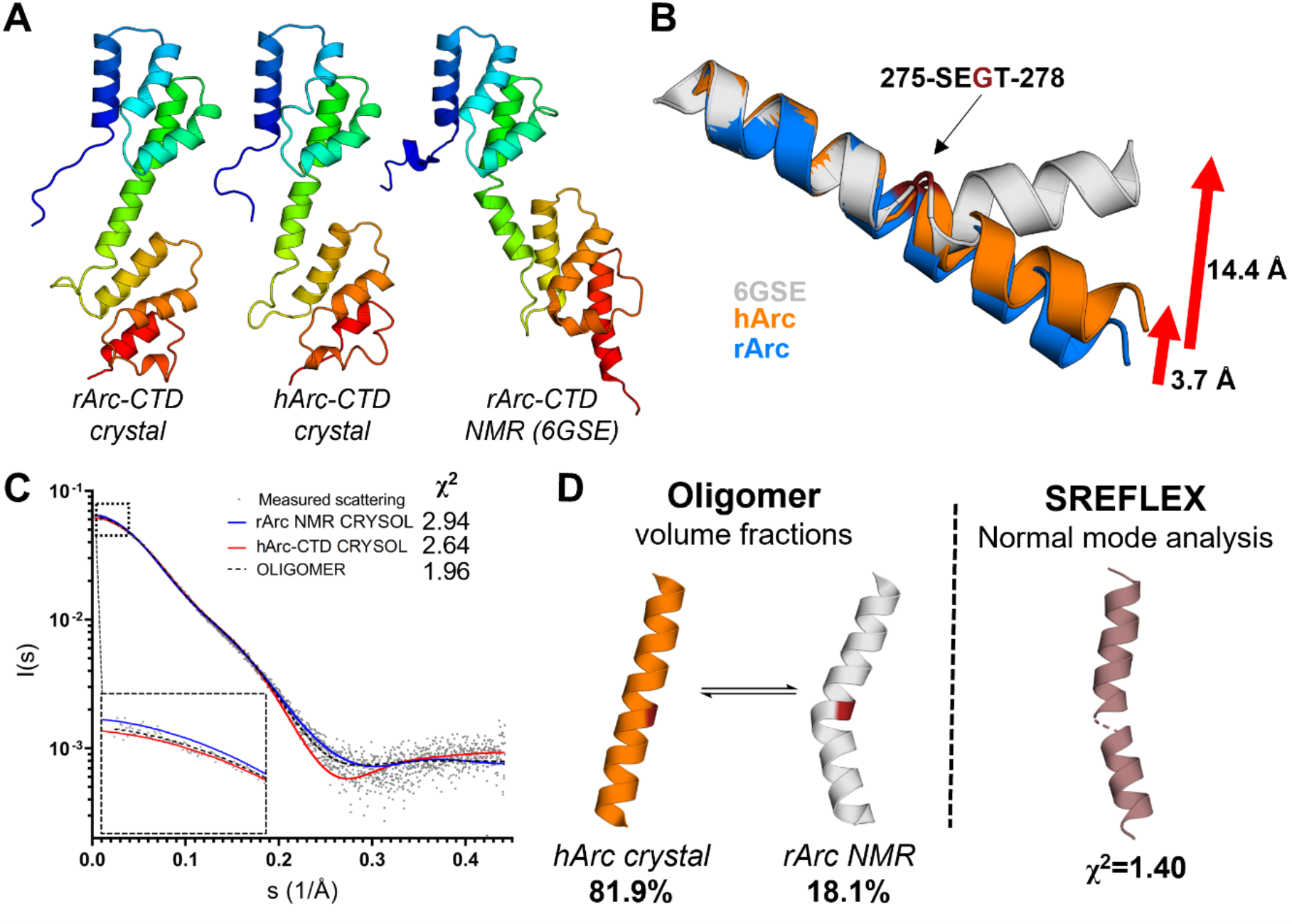
The Arc-CTD is dynamic in structure. **A** Comparison of the two Arc-CTD crystal structures with the NMR model of rArc-CTD (6GSE, [22]) reveals a hinge region in the central helix. **B** A closer look at the helices, superimposed on the N-terminal end, highlights a hinge region centred on Gly277. **C** Analysis of the solution structure of the Arc-CTD using SAXS. Shown is the experimental scattering curve of the HArc-CTD construct, theoretical scattering curves generated using CRYSOL for the crystalline and the previously published NMR model (6GSE, [22]), and the fit from volume fraction analysis of the two in OLIGOMER. **D** The orientation of the two helices and results of volume fraction analysis (left). Normal mode analysis (right) of the hArc-CTD crystal structure against the SAXS gave the most plausible average solution conformer of the central helix. The N- and C-lobes are not shown in panels C and D for clarity.

To estimate the conformational flexibility of the CTD in solution, SAXS was used (Fig. 10C-D). Calculation of theoretical scattering curves from the hArc-CTD crystal structure and the NMR model of the rArc-CTD and fitting against SAXS data, previously obtained from hArc-CTD without the N-terminal His6-tag [26], showed that neither seemed representative of the observed solution scattering. Analysis of the representative volume fraction of each conformer gave fractions of around 80% and 20% for the crystal and NMR conformations, respectively. This suggested that the mean solution conformer was closer to what was observed in the crystal structure, but not identical. Furthermore, normal mode analysis of the hArc-CTD crystal structure against the SAXS data gave an average conformer that showed a slightly kinked helix closer to the crystallised conformation. Thus, considerable conformational flexibility was present in the CTD, which has a dynamic structure.

To obtain a more detailed insight into conformational plasticity of the CTD, the hArc-CTD crystal structure, stripped of bound Nbs, was subjected to a 640-ns atomistic MD simulation (Fig. 11). In the simulation, the breaking of the central helix at Gly277 facilitates extension of the central hinge region, and after rotation of the two lobes they pack onto each other, resulting in a complete collapse after only 50 ns. This collapsed conformation seemed more stable than the extended conformer, as little conformational change was observed for the remainder of the simulation. Therefore, the simulation further suggested the CTD to be dynamic in structure and implied that hArc-CTD can fold into conformers similar to that of the capsid protomers of dArc1 and dArc2.

**Figure 11:**
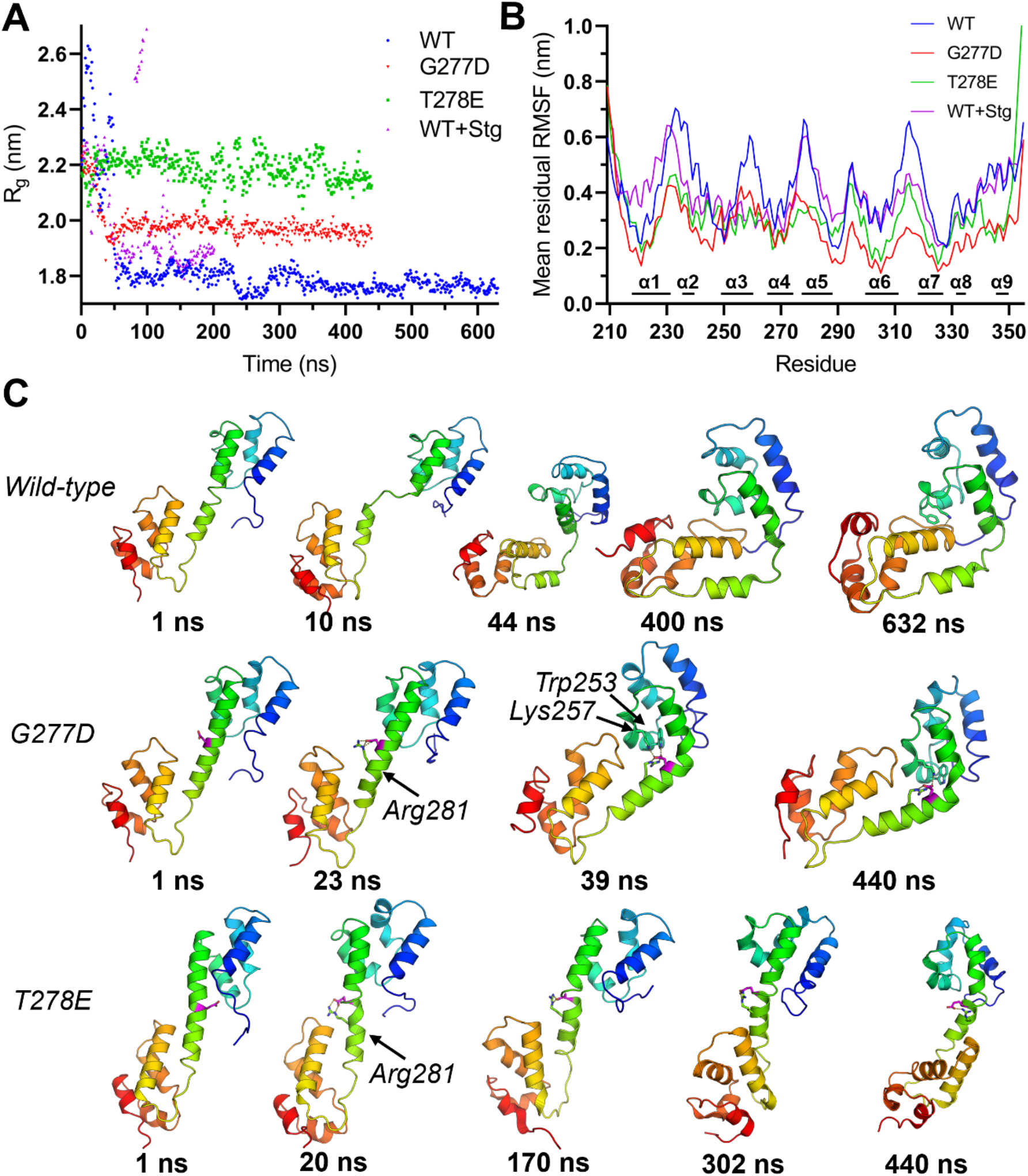
MD simulations of wild-type (WT) CTD with and without bound Stg and mutants G277D and T278E. **A** The calculated R_g_ over the course of the trajectories clearly demonstrate the compaction of the wild-type and its reduction in the G277D and T278E mutants. **B** Mean residual RMSF for all residues demonstrates the reduced flexibility of the mutants. **C** Trajectory snapshots at selected time points. The *in silico* mutated residues in G277D and T278E are highlighted in magenta.

Thr278 of the Arc-CTD undergoes phosphorylation by TNIK kinase *in vivo*, reducing the ability of fulllength Arc to form capsids [69]. Furthermore, the G277D mutation has been suggested to reduce the ability of Arc to oligomerise [36]. As both mutations are located in the dynamic hinge region of the CTD central helix, MD simulations were run for G277D and T278E, to examine if altered dynamics would be observed in these oligomerisation-inhibiting mutants (Fig. 11). In the G277D simulation, the helix did break but complete compaction of the domain was restricted by the negative charge introduced by Asp277. In the case of the T278E phosphomimic, as Glu278 formed a salt-bridge with the adjacent Arg281 to restrict the initial extension of the hinge region, inhibiting the collapse of the domain. As these mutations reduce the ability of Arc to oligomerise, the simulations suggest a role for the CTD structural dynamics in Arc capsid assembly.

### Crystal structure of a collapsed Arc-CTD

Considering the dynamic nature of the Arc-CTD, crystallisation of the protein in different conformers was attempted. Optimisation of previously neglected crystallisation conditions for hArc-CTD in complex with NbArc-H11 and -C11 was carried out. Initial optimisation slightly improved crystal quality, and matrix microseeding produced crystals of sufficient quality for structure determination. These crystals proved to be of higher quality than those used to solve the structures of rArc- and hArc-CTD, allowing for structure determination at 1.94-Å resolution (Fig. 12).

**Figure 12:**
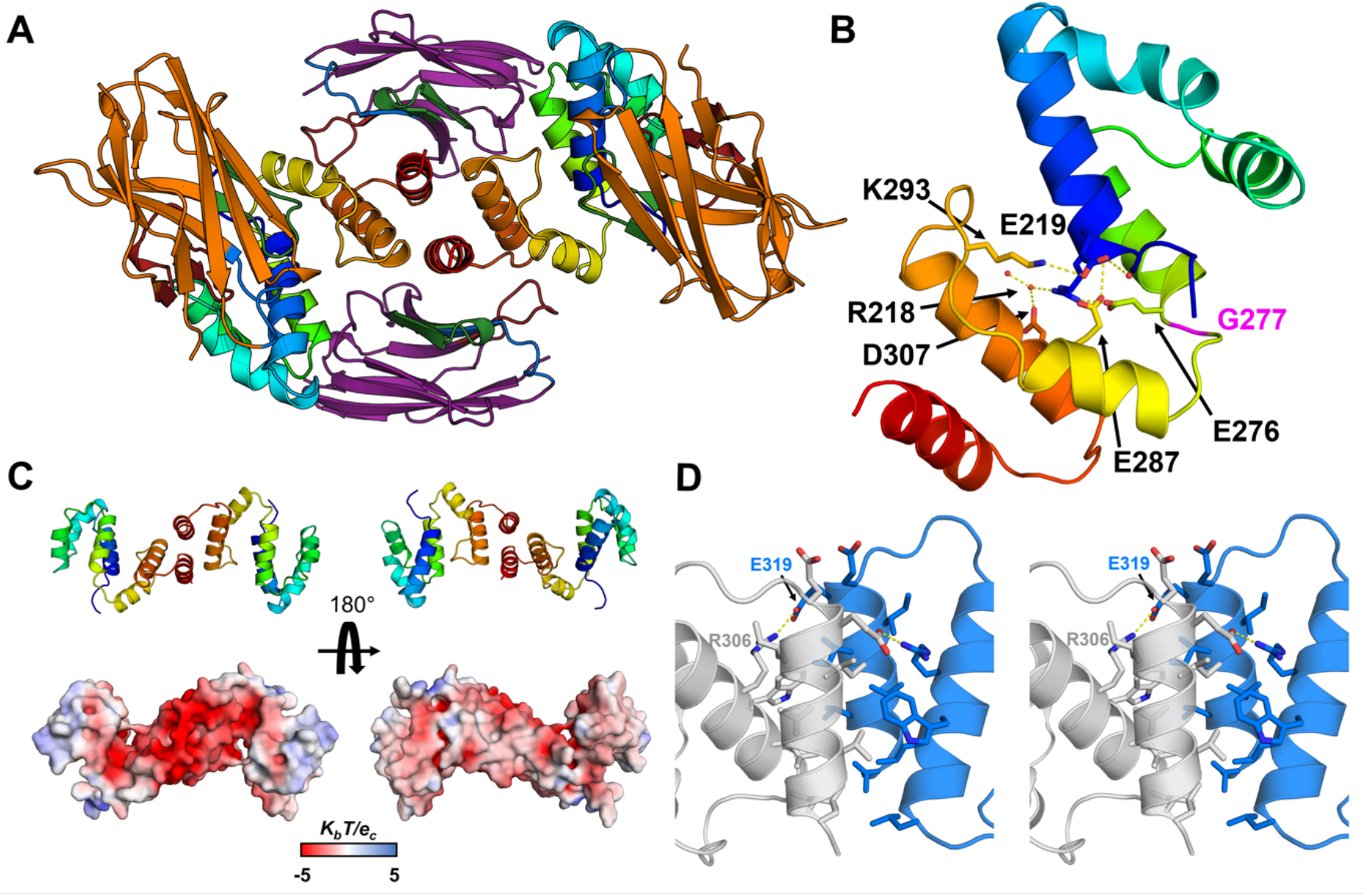
1.94 Å crystal structure of the dimeric broken helix CTD of hArc in complex with NbArc-H11 and -C11. **A** The overall structure of the dimeric hArc-CTD+H11+C11 complex observed in the crystal, viewed down the dimer symmetry axis. The two CTDs are shown in rainbow, H11 in orange and C11 in purple. CDR1, 2, and 3 are coloured blue, green and red, respectively. **B** A single monomer, highlighting the collapse of the domain and lack of the last two helices of the C-lobe. Shown are the sidechains which facilitate interactions on the hydrated inter-lobal interface. Gly277, of the central helix hinge region, is shown in magenta. **C** Dimerisation results in rearrangement of charges, resulting in one highly acidic surface side. **D** The putative CTD dimer interface observed in the crystal (stereo view); the two monomers are coloured blue and grey. The ten residues of each monomer that form the interface are shown, and the inter-monomer salt bridge formed between Glu319 and Arg306 is highlighted.

The complex crystallised in a different unit cell, when compared to the two earlier structures of the rat and human CTD. Accordingly, the structure of the complex was altered (Fig. 12A). In this conformer, breaking of the central helix at Gly277 leads to a complete collapse of the CTD and the packing of the C-lobe against the N-lobe (Fig. 12B). The interface between the lobes remained partially hydrated and was mainly characterised by indirect hydrogen bonding or inter-lobal salt bridges, such as those involving Glu219, Glu276, Glu287, Arg218 and Lys293. Moreover, the CTD appeared dimeric. Using PISA, the calculated free energy gain upon dimerisation (ΔG^int^) of −54.4 kJ mol^−1^ indicated a stable dimeric assembly. The putative dimer interface (Fig. 12D), which buries 600 Å^2^ of surface area from each monomer, formed between the C-lobes and involves 10 residues of each subunit. Residues 212-330 of CTD could be built into the electron density, but the last two helices of the domain were missing. Therefore, the dimer interface formed between incomplete C-lobes, fusing the exposed hydrophobic cores of the two protomers. Accordingly, the dimer interface is characterised by non-polar interactions, except for a salt bridge formed between Glu319 and Arg129. What led to the absence of the C-terminal end of the CTD, and dimerisation, is unresolved. Either the missing C-terminal portion was lost due to degradation, or it was not observed due to flexibility. In support of the latter, large empty spaces were observed in the crystal lattice in proximity of the C-termini (Fig. S4C). Interestingly, the observed dimerisation scheme led to a rearrangement of the electrostatic surface potential of the CTD, resulting in one side of the dimer being highly acidic while the other, with both C-termini, has a more neutral surface potential (Fig. 12C). This might facilitate interactions with cationic components, such as the Arc NTD. The unique conformer observed in the crystal resembles that observed in MD simulations and could be representative of the hArc capsid protomer, as the fold shares high resemblance with a variety of viral capsid protomers, including dArc1 and dArc2.

With the ability of Nbs to bind and select for unique protein conformers, it was surprising that both NbArc-H11 and -C11 bound to the collapsed conformer (Fig. 12A). In particular, a portion of the C11 epitope, present on the last two helices of the C-lobe was absent in the structure, and the surface area C11 buries from the C-lobe was reduced to 658 Å^2^. This was seemingly compensated by additional interactions of the FR of C11 with the N-lobe of the CTD, not observed in the other crystal structures, which account for an additional interacting surface of 271 Å^2^. Whether these additional interactions are representative of binding interactions in solution or arise simply due to crystal packing is uncertain. In addition, the binding of H11 did not restrict the conformational shift. Upon the collapse of the CTD, the second loop of the C-lobe (the *α*5-*α*6 loop of the CTD) comes into proximity of the CDR2 loop of H11, showing modest interactions but no apparent steric hindrance. Otherwise, the overall binding mode of H11 was nearly identical to what was observed in the other crystal structures.

### Solution structures of hArc-CTD-Nb complexes

Considering the lack of strict conformational selection by NbArc-H11 and -C11, as they crystallised in complex with both extended and collapsed forms of the CTD, the solution structures of all Nbs in complex with Arc CTD were investigated using SEC-SAXS (Fig. 13, Table 5). The aim was to determine whether conformational selection occurred and allow for coarse epitope mapping. In all cases, the Nbs bound the monomeric hArc-CTD in 1:1 stoichiometry, suggesting that the dimeric hArc-CTD+H11+C11 crystal structure was either an artefact of crystallisation, or that it only occurred following the degradation or unfolding of the C-terminal end of the CTD.

**Figure 13:**
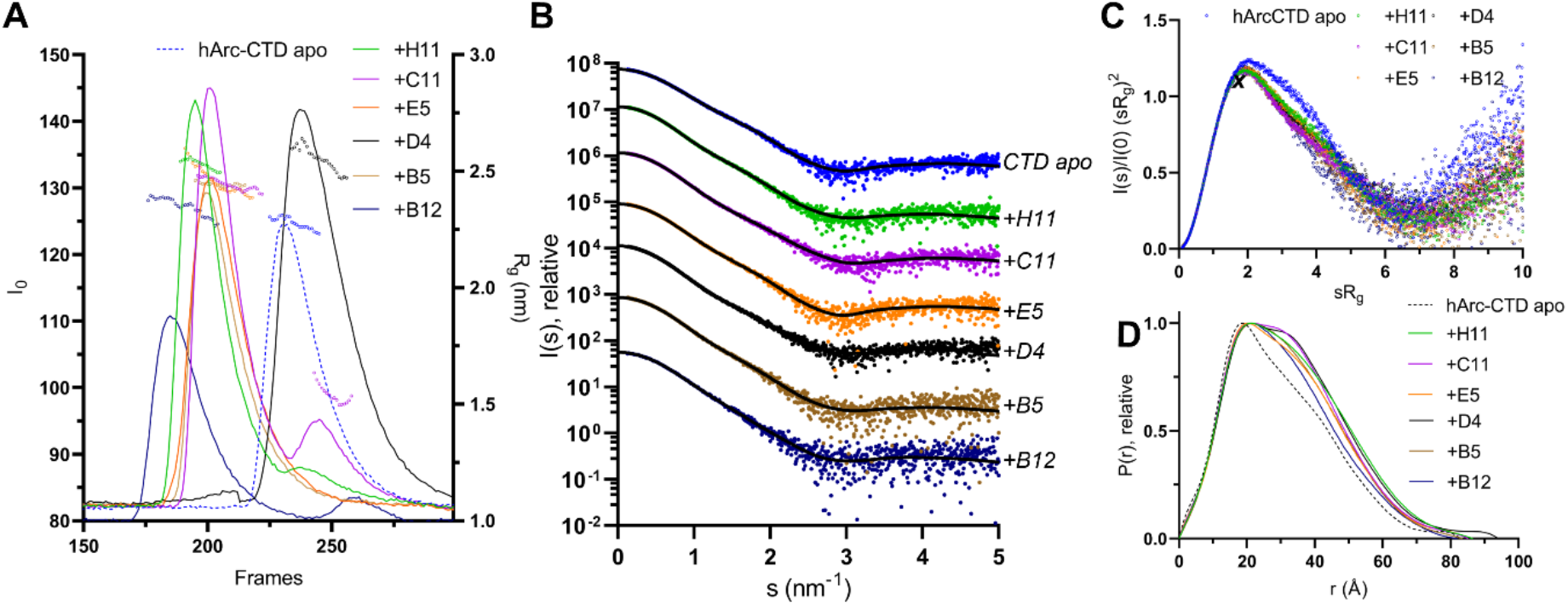
SEC-SAXS of hArc-CTD Nb complexes. **A** SEC-SAXS elution profiles. The observed scattering intensity is shown as lines (left Y-axis) and the calculated R_g_ is shown for the frames used for data processing. The R_g_ of the Nb excess peaks is shown for C11 and H11+C11 complex run, although they were not used for processing. **B** 2D scattering curves obtained from the main SEC peaks. Curves are offset by one logarithmic unit, for clarity, and data fits from GNOM shown as unbroken lines. **C** Dimensionless Kratky plot. The maximum for an ideal rigid spherical particle is marked with ***X*** (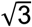, 1.104). **D** Distance distribution profiles. The free hArc-CTD (apo) profile is shown as black dashes.

**Table 5:** Size and shape parameters for CTD-Nb complexes, derived from the SAXS data in Fig. 13.

| Protein | hArc-CTD | +H11 | +C11 | +E5 | +D4 | +B5 | +B12 |
| --- | --- | --- | --- | --- | --- | --- | --- |
| $I_0$ | 74.85 | 114.99 | 116.33 | 90.89 | 112.38 | 86.02 | 55.87 |
| $R_g$ (Å) | 23.0 | 25.3 | 24.6 | 24.5 | 25.5 | 24.4 | 23.7 |
| Porod volume (nm<sup>3</sup>) | 29.22 | 43.02 | 40.37 | 40.44 | 42.51 | 42.07 | 38.93 |
| $P(r) R_g$ (Å) | 23.4 | 25.7 | 25.0 | 24.9 | 26.0 | 24.7 | 24.2 |
| $D_{max}$ (Å) | 83.0 | 86.5 | 85.2 | 85.1 | 94.0 | 83.5 | 82.0 |
| Quality est. GNOM | 0.71 | 0.87 | 0.83 | 0.80 | 0.72 | 0.87 | 0.85 |
| $Q_p$ Mw (kDa) | 21.4 | 32.1 | 29.9 | 30.1 | 30.6 | 31.5 | 28.8 |
| Monomer mass (kDa) | 21.7 | 35.8 | 34.9 | 34.5 | 34.6 | 34.9 | 34.4 |
| Oligomeric state/Stoichiometry | Monomeric | 1:1 | 1:1 | 1:1 | 1:1 | 1:1 | 1:1 |

In complex with the Nbs, the CTD seemed to compact to some extent, as the Kratky plots (Fig. 13C) suggested that the complexes were more rigid and compact than the free CTD. This was also apparent in the obtained size parameters. R_g_ only slightly increased and D_max_ remained almost unchanged despite the increased Porod volume and molecular weight, which both increase by ~50% upon Nb binding (Table 5). This was observed most clearly for the B12 complex. As the Nbs approximate the hArc-CTD in mass, one would have expected a greater size increase in the complexes. Compactness was further highlighted in the calculated distance distribution of the complexes (Fig. 13D), where the split profile in the unbound protein, characteristic of a bilobar structure, is lost into a single wider peak, accompanied by little change in D_max_. The CTD compaction upon Nb binding also seems apparent in the *ab initio* models of the complexes (Fig. S5). The models of the complexes were similar in size to the free CTD, and the linker in the apo domain was not as apparent. Although epitopes could not be mapped with SAXS due to the symmetric nature of the CTD, it seemed that all Nbs associated mostly with either the N- or the C-lobe. Altogether, the SAXS data suggested that Nb binding, to either of the two lobes, drives compaction of the CTD *via* conformational selection. Compaction of the CTD in full-length Arc might be facilitated by interdomain interactions with the NTD.

### The coiled-coil NTD mediates dimerisation of FLrArc-7A, indicating a role in higher-order assembly

To assess the role of the individual domains of the FLrArc-7A mutant in dimerisation and higher-order oligomerisation, solution structures and oligomeric states were assessed using SAXS. The Arc-NTD is poorly soluble, and the isolated domain resisted recombinant expression and purification (data not shown). Therefore, the rArc-NTD with the oligomerisation-inhibiting poly-Ala mutation was expressed and purified as a soluble MBP fusion (MBP-2rNT), and SAXS was used to study solution structure and oligomeric state (Table 6, Fig. S6). Solution scattering data showed that MBP-2rNT was dimeric and rigid, in contrast to the monomeric hArc-CTD (Table 5). This indicated that dimerisation of the full-length FLrArc-7A is likely mediated by dimerisation of the NTD. The *ab initio* model of MBP-2rNT (Fig. S6) revealed a rod-shaped fold with lobes on opposing ends, which represent the N-terminal MBP fusions at opposing N-termini. The density at the centre of the envelope likely accounts for a four-helix bundle formed by end-to-end packing coiled coils. This hypothetical model of MBP-2rNT was further explored with *in silico* modelling in CORAL, using the previously published homology model of the NTD [26] and a crystal structure of MBP (Fig. 14). In this conformation, the two oligomerisation motifs, ^113^MHVWREV^119^, are located at opposite ends of the four-helix bundle. This, together with the dynamic nature of the CTD, might provide insights into the mechanism of Arc oligomerisation.

**Table 6:** SAXS analysis of the Arc NTD.

| Protein | MBP-2rNT |
| --- | --- |
| I<sub>0</sub> | 0.053 |
| R<sub>g</sub> (Å) | 38.79 |
| Porod volume (nm<sup>3</sup>) | 155.0 |
| P(r) R<sub>g</sub> (Å) | 39.91 |
| D<sub>max</sub> (Å) | 125.15 |
| Quality est. GNOM | 0.90 |
| Q<sub>p</sub> Mw (kDa) | 109.9 |
| Monomer mass (kDa) | 56.0 |
| Oligomeric state | Dimer |

**Figure 14:**
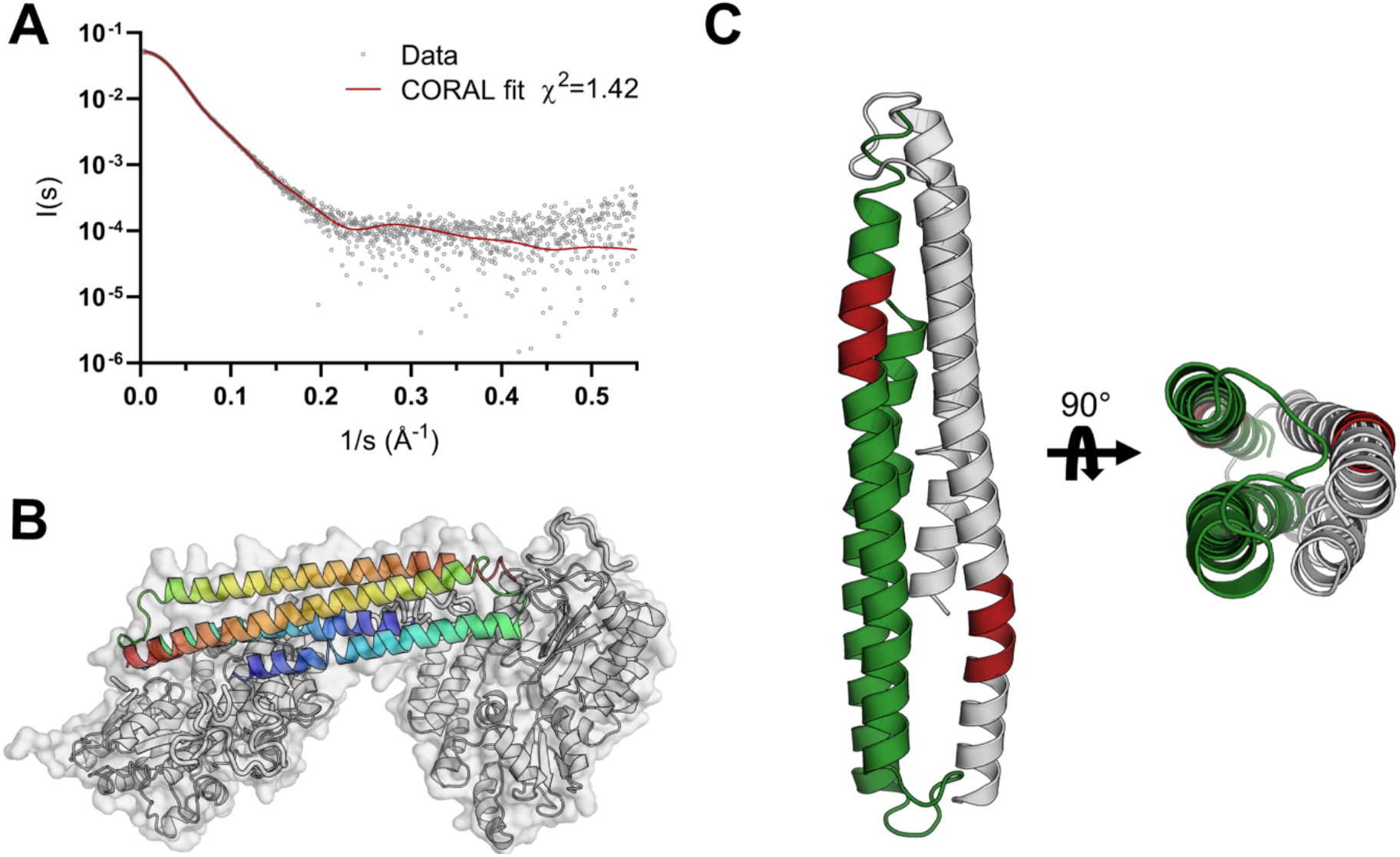
*In silico* model of the rArc-NTD, based on homology modelling and SAXS data. **A** The data fit obtained from CORAL, indicating a good fit to the data. **B** The hypothesised low-resolution model of the MBP-2rNT dimer, obtained from the data fit shown in a). The model was produced using a homology model of the wild-type NTD [26] and a crystal structure of MBP (1NL5, [89]). The two coiled coils are coloured blue to red from N- to C-termini, respectively, and the two MBPs in grey. **C** The arrangement of the NTD in the model, showing a four-helix bundle consisting of antiparallel coiled coils. The position of the oligomerisation motif (mutated in this construct) is shown in red.

## Discussion

In this study, we characterised six ultra-high affinity anti-Arc Nbs. Two of the Nbs, NbArc-H11 and -C11 were used to obtain crystal structures of the rat and human Arc-CTD. The crystallisation of the hArc-CTD in two different conformations provided an insight into the structural dynamics of the domain, which was further explored using SAXS and MD simulations, suggesting an important role of these dynamics in higher order oligomerisation, and capsid formation.

### The CTD conformational change and its physiological relevance

The hArc-CTD crystallised in the extended conformation at a high concentration (37 mg/mL) with Nbs H11 and C11, suggesting a solubilising effect by Nb binding. However, at a lower concentration (10 mg/mL), the complex crystallised in a collapsed conformer. What led to the crystallisation of this unique conformer of the hArc-CTD, is uncertain. Possibly, the low ionic strength of the crystallisation condition had an effect; capsid formation by Arc is dependent on ionic strength [32]. A more likely explanation lies in the absence of the C-terminal end of the CTD in electron density. Unfolding or degradation of the last two helices of the hArc-CTD resulted in dimerisation, in which the exposed hydrophobic cores of the remaining C-lobes formed the dimer interface. Interestingly, this dimerisation mode resulted in one highly acidic face, while both C-termini localised on the opposite side. In the viral-like capsids formed by dArc1, the 48-residue C-terminal tail is located inside of the capsid. Two tails of the hexameric capsomer subsequently form a two-stranded zinc finger to facilitate RNA binding, while the remaining four remain unstructured [34]. Similarly, the collapsed conformer of the hArc-CTD could represent the structure of the CTD in hArc capsids, whereby the acidic side binds the basic NTD on the outer capsid layer, while the C-terminal tail locates to the inside. Moreover, phosphorylation sites have been identified in the in unstructured C-terminal portion of Arc, at residues Thr368 and Thr380, the phosphorylation of which regulates the degradation of Arc [70, 71]. Whether these post-translational modofications facilitate the unfolding of the C-terminal end of the CTD and the subsequent conformational shift and dimerisation, and possibly the higher-order oligomerisation of Arc, remains a subject of further study.

### Peptide binding displacement and its effects on Arc structure and function

The Arc-CTD binds ligand peptides in a hydrophobic groove of the N-lobe. The CDR3 of NbArc-H11 contained the conserved residues of the Arc ligand peptide motif and bound in the same binding of the CTD co-crystal structures. We further demonstrated, *via* an ITC displacement assay that H11 could efficiently displace the Stg peptide bound to the FLrArc-7A CTD. In the structure of the hArc-CTD N-lobe in complex with the Stg peptide, the portion of the NTD-CTD linker preceding the CTD N-terminus folds against the peptide as it bound in the hydrophobic groove of the N-lobe [24]. In the structure of the hArc-CTD, obtained here *via* co-crystallisation with NbArc-H11 and -C11, the same segment extends away from the N-lobe (Fig. 15A). These findings propose that the binding of Stg can alter the structure of the NTD-CTD linker region, resulting in large changes in the relative orientation of the two folded domains of Arc (Fig. 15B). This could represent a previously unrecognised mechanism of how Stg, and presumably other Arc ligand peptides, might regulate the interdomain interactions of full-length Arc and possibly higher-order oligomerisation.

**Figure 15:**
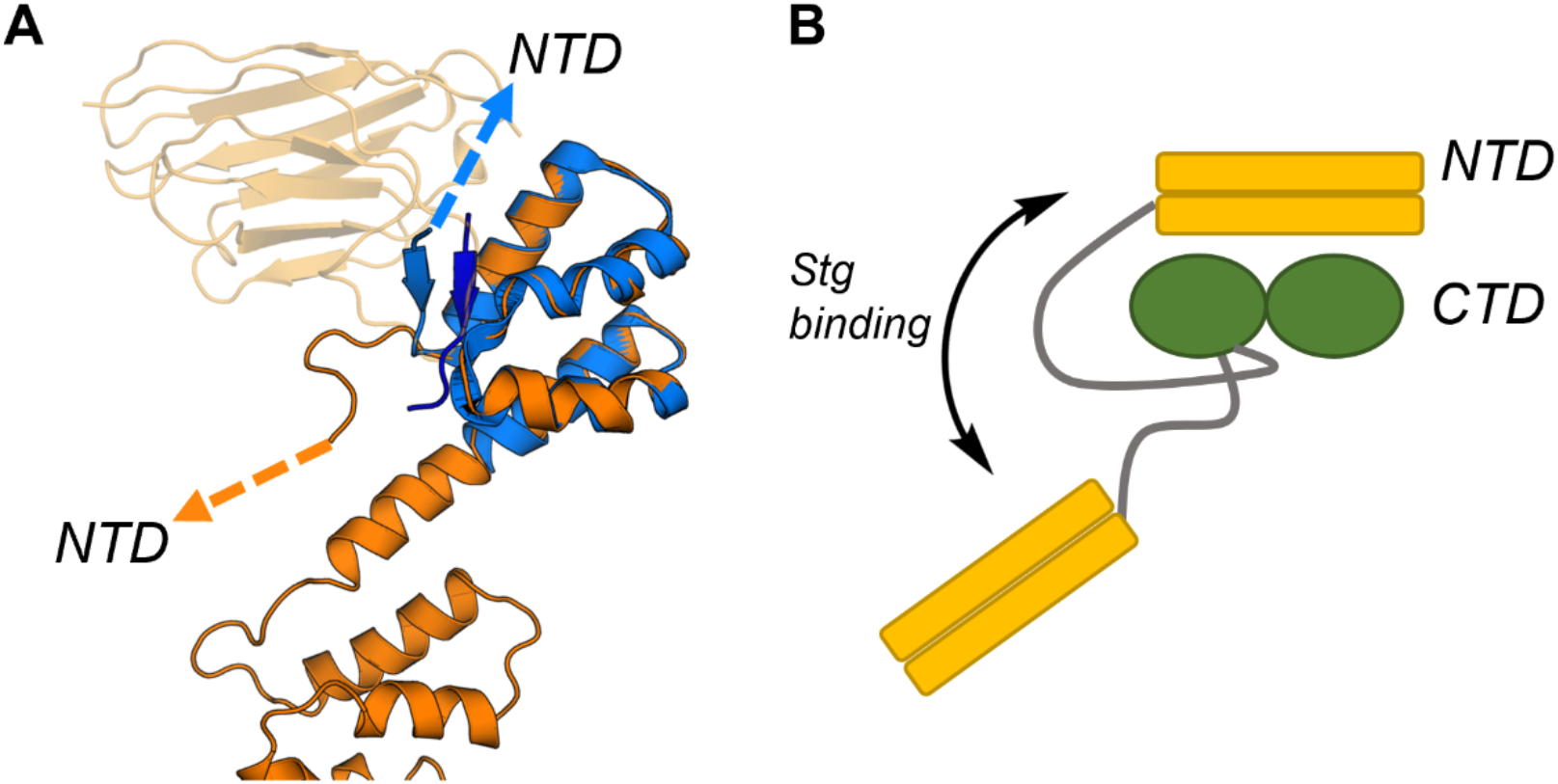
The potential effect of Stg binding on the structure of full-length Arc. **A** Alignment of the extended conformer of the hArc-CTD (orange), crystallised in complex with NbArc-H11 and -C11, and the isolated hArc-CTD N-lobe crystallised in complex with Stg (blue, 6TNO, [24]). NbArc-H11 is shown in transparent orange and the bound Stg in dark blue. The conformational change in the linker region is noted, by arrows pointing towards the NTD. **B** The hypothesised effect of Stg binding on the structure of linker region and the relative orientation of the two domains of Arc. The linker region is shown in grey.

Nbs have in recent years emerged as powerful tools for probing protein structure, function, and dynamics *in vivo*. The monomeric and soluble nature of Nbs allows for, in many cases, their correct folding and function in the reducing environment of the eukaryotic cytosol. Moreover, as they only consist of a single domain, the manipulation of their sequence and addition of various fusion proteins is made possible. As an example, transgenic expression of a green fluorescent protein (GFP) fused Nb, a so called chromobody, can allow for real-time visualisation of subcellular localisation of the antigen in live cells [72], and co-expression of Nbs fused to subunits of the E3-ubiquitin ligase complex rapidly facilitates ubiquitinoylation and degradation of the target protein [73, 74]. In addition, the conformationally selective nature of Nb binding has been utilised to stabilise transient, physiologically relevant conformers and to probe conformational dynamics *in vitro* and *in vivo* [75, 76]. Arc has several putative demonstrated interaction partners in the PSD, including Stg, Glu2NA, GKAP, as demonstrated by binding of ligand peptides in the hydrophobic groove of the CTD N-lobe [18, 22, 24]. The physiological relevance of these interactions, however, remains poorly understood. The H11 Nb characterised here presents itself as a highly useful tool for studying these interactions as it was shown to efficiently displace bound peptide without affecting other Arc structural integrity or noticeably affecting capsid formation in purified Arc.

In parallel with the present work, we have generated recombinant epitope-tagged Nbs for immunoblotting and affinity-purification of endogenous Arc from brain tissue samples, including Arc expressed after induction of synaptic plasticity (long-term potentiation) in live rats (Ishizuka et al., in preparation). Epitope mapping, based on expression of Arc regions in cell lines, shows that H11 and E5 selectively bind the N-lobe, while B5, B12, C11, and D4 bind the C-lobe. When expressed as genetically encoded intrabodies, H11 and E5 bound to the N-lobe and allowed immunoprecipitation of full-length Arc. In addition, Nbs fused to the fluorescent mScarlet expressed as chromobodies (and localized to induced endogenous Arc). We therefore foresee many applications of Arc Nbs as modular tools for studying Arc function, subcellular localisation, and structural dynamics *in vivo*.

### Conformational flexibility of the CTD and dimerisation of the NTD, and their potential roles in capsid formation of Arc

Using NbArc-H11 and -C11 as crystallisation chaperones, the CTD of rArc could be crystallised. Moreover, the same Nbs allowed for the crystallisation of the hArc-CTD, which prior to this project had not been structurally characterised, in both an extended and a collapsed conformation, painting a highly dynamic picture of the domain. The dynamic nature of the CTD was further investigated using SAXS and MD simulations. In MD simulations of the G277D and T278E mutants, this conformational plasticity was lost and the CTD remained in more extended conformers. As these mutations, and especially the T278E phosphomimic, have been reported to reduce the ability of full-length mArc to form viral-like capsids [36, 69], the simulations suggested a fundamental role of the structural plasticity of the CTD, and the collapsed form observed in the crystal structure, in capsid formation.

The fold similarity of the N- and C-lobes of dArc and mArc-CTD with various retroviral Gag polyprotein capsid (CA) domains has been demonstrated [18, 24, 30, 35]. In the dArc capsids, and the capsid formed by the homologous Ty3/Gypsy retrotransposon [34, 77], the two lobes are connected by a short linker, similar to that observed in the collapsed crystal form of hArc-CTD (Fig. 16A). From the capsid structures, it seems that breaking of the CTD linker region is needed for capsid formation. Similarly to Arc and Ty3/Gypsy, the CA domain of the HIV Gag polyprotein contains two lobes connected by a short linker. Flexibility of this linker is vital HIV for capsid assembly [78, 79]. Moreover, comparison of the monomeric HIV CA domain crystallised in complex with a Fab fragment [80] and the pentameric capsid protomer reveals a conformational change, similar to that observed for Arc-CTD, between the monomeric and capsid forms (Fig. 16B). The same applies to Rous Sarcoma Virus (RSV), where an extended conformer is found in immature viral particles, and movements in the hinge region facilitate formation of mature RSV capsids [81, 82] (Fig. 16C). These similarities suggest that the structural plasticity of the Arc-CTD might represent a conserved mechanism for capsid assembly of retroviruses and LTR retrotransposons.

**Figure 16:**
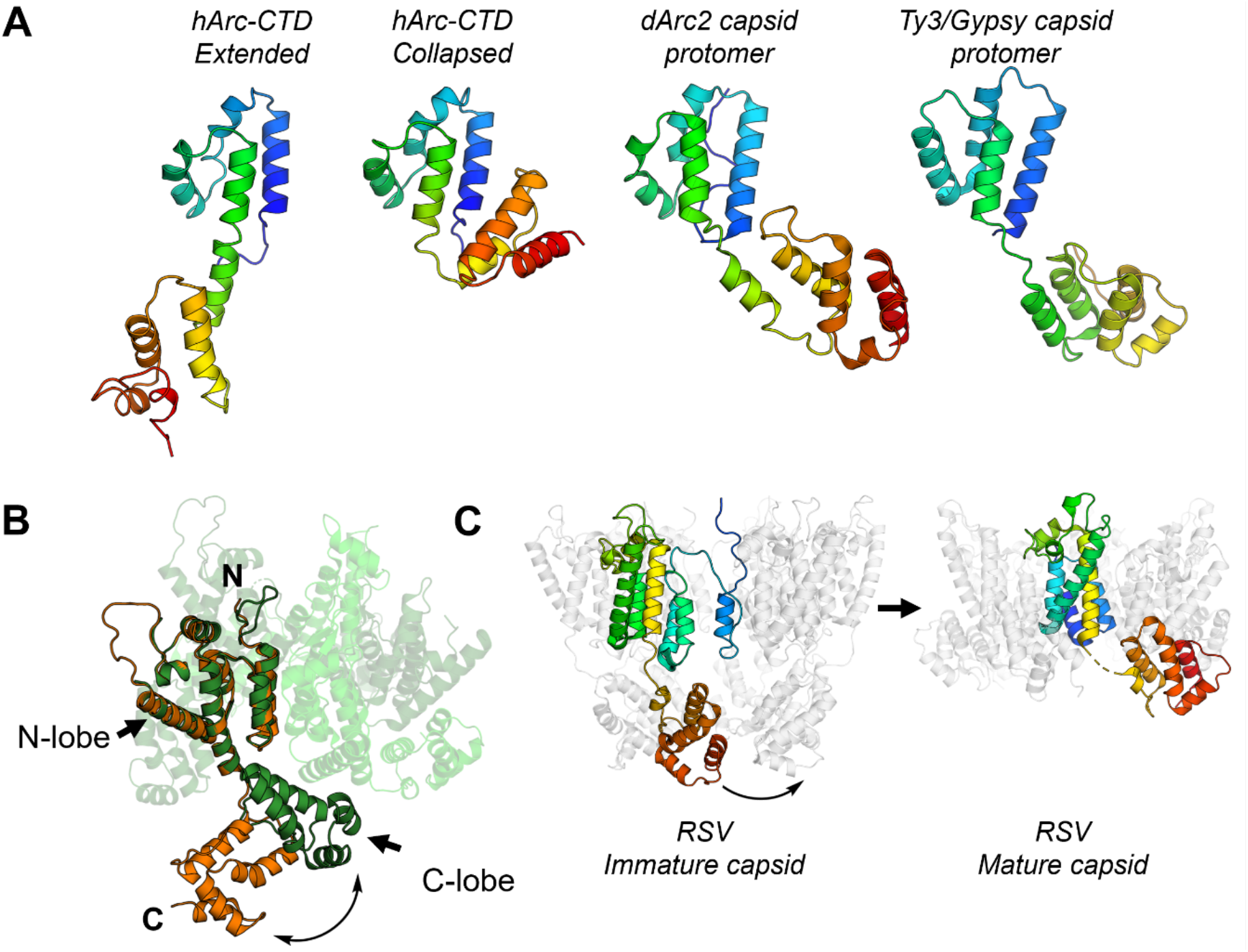
CTD hinge region flexibility is conserved in other LTR retrotransposons and retroviral CA domains. **A** Comparison of the extended and collapsed crystal structures of the hArc-CTD, obtained in this study, with the dArc2 (6TAQ, [34]) and Ty3/Gypsy (6R24, [77]) capsid monomer reveal varying conformations of LTR retrotransposon capsid proteins. All are aligned on the N-lobe and shown in rainbow. **B** Comparison of the monomeric HIV CA, crystallised in complex with a Fab fragment (orange, 1E6J, [80]), and the CA in the capsid pentamer (3P05, [78]) reveal structural plasticity of the linker region, similar to that observed in the Arc-CTD. The HIV CA monomer is shown in orange and the capsid pentamer in green. The capsid protomers not aligned with the monomeric CA are shown in various shades of green transparent cartoons. **C** Structure of immature RSV viral particles (5A9E, [81]) and mature RSV capsid pentamers (7NO5, [82]) shows similar structural fluctuations in the process of capsid assembly.

The viral-like capsids of dArc1 and dArc2 consist of penta- and hexameric capsomers, the formation of which depends on homo-oligomerisation of the N-lobe. Consequently, assembly of the mature capsid is facilitated by dimerisation of the C-lobe [34]. This is expedited by the strong tendency of the individual lobes of both dArc isoforms to homo-oligomerise [35]. In contrast, the CTD of mArc is monomeric in solution, as observed here for hArc and as previously been demonstrated [26]. Importantly, a determining factor of mArc capsid assembly is the oligomerisation motif in the second coil of the coiled-coil NTD [33], which was mutated in the dimeric FLrArc-7A construct used here. Nevertheless, the NTD does not independently assemble into capsids in the absence of the CTD, suggesting a role for both domains in capsid assembly [32]. Previous studies have hypothesised on the determinants of hArc capsid assembly, suggesting domain swapping of the NTD and CTD, as two rigid bodies, to be the determining factor of oligomerisation [26]. However, the structural plasticity of hArc-CTD and its similarity to that of other retrotransposons and retroviral capsid proteins (Fig. 16) suggest that the hArc-CTD might form penta- and hexameric capsomers in an icosahedral capsid, similarly to dArc. Moreover, solution scattering of the soluble MBP-fused, mutated NTD construct showed the rArc-NTD to be homodimeric. In the suggested model of the NTD (Fig. 14), the oligomerisation motifs are situated at opposing ends of the dimer.

Based on the current data, a model for the hArc capsids can be speculated upon, in which the dimerisation and further oligomerisation of the NTD coil motif facilitate capsomer formation and link adjacent capsomers. The capsomeric CTD may be similar to the collapsed hArc-CTD crystal structure, and additional dimerisation of the CTD C-lobe cannot be excluded. The NTD has been demonstrated to bind and induce curvature in anionic membranes [26, 27], and would therefore likely be situated on the outside of the capsid to facilitate intercellular transport. While other models of capsid assembly cannot be excluded, this hypothesis explains the dependency of hArc on the NTD oligomerisation motif and accounts for both NTD dimerisation and CTD structural plasticity. Dimerisation *via* the NTD could be achieved through domain swapping, in agreement with earlier hypotheses [26], which would approximate the inter-capsomer domain swapping observed in Simian virus 40 and polyomavirus capsids [83, 84]. In this model, the 84-residue linker connecting the NTD and CTD would have to traverse from the centre of the capsomer towards the dimerising NTDs. The average end-to-end distance of an 84-residue disordered chain is around 8.3 nm [85], and with a typical *C_α_-C_α_* distance of 3.8 Å, the length of the linker could, theoretically, extend up to ~30 nm. Its length should, therefore, not be a limiting factor in the model. The validation of such a model remains a subject of further study.

### Concluding remarks

We have presented a series of recombinant ultrahigh-affinity nanobodies directed against the N- and C-lobes of the mammalian Arc CTD. As shown here, these nanobodies are functional tools for structural biology approaches, and they can be further developed towards applications in *e.g*. imaging and functional modulation of Arc. From the structural studies enabled by the nanobodies, we have obtained crystal structures of the Arc CTD in two different conformations, highlighting a hinge region in the linker helix between the CTD lobe domains as possibly important for Arc functional conformational changes and virus-like capsid formation. Future work using these nanobody tools will enable novel, exciting research avenues in structural and functional biology of mammalian Arc.

## Acknowledgements

We acknowledge the use of the Core Facility for Biophysics, Structural Biology and Screening (BiSS) at the University of Bergen, which has received funding from the Research Council of Norway (RCN) through the NORCRYST (grant number 245828) infrastructure consortium. Electron microscopy was carried out at the Molecular Imaging Centre (MIC) at the University of Bergen. This work was supported by a Research Council of Norway TOPPFORSK grant (249951) to CRB.

Parts of this research were carried out on beamline P11 at DESY, a member of the Helmholtz Association (HGF). We wish to thank EMBL/DESY for access to beamline P14 and acknowledge Diamond Light Source for time on Beamlines I03 under Proposal MX18666. In addition, beamline support at SWING (SOLEIL synchrotron) and BM29 (ESRF) is acknowledged. The simulations were performed on resources provided by UNINETT Sigma2 - the National Infrastructure for High Performance Computing and Data Storage in Norway, on the Fram and Saga computing clusters under project number NN9859K.

## Notes

### Competing Interest Statement

The authors have declared no competing interest.

### Summary of Updates

Added one author, who accidentally was missing from the form.

