## Supplementary Information for "High-affinity nanobodies as tools for structural and functional studies on mammalian Arc"

### Supplementary Figures S1-S5

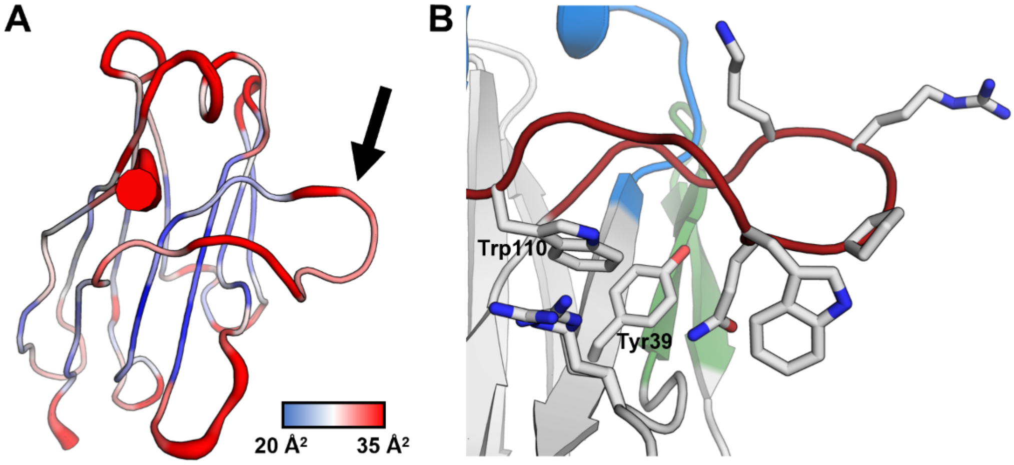


**Figure S1:** Flexibility of the CDR loops of NbArc-E5. **A** The crystallographic B-factors of the NbArc-E5 crystal structure demonstrate the rigidity of the CDR3 loop (highlighted by arrow). **B** Rigidity of the CDR3 loop is likely accounted for by packing of its large hydrophobic side chains onto the exterior of the β-barrel fold. This could also account for the increased solubility of NbArc-E5 when compared to the other Nbs.

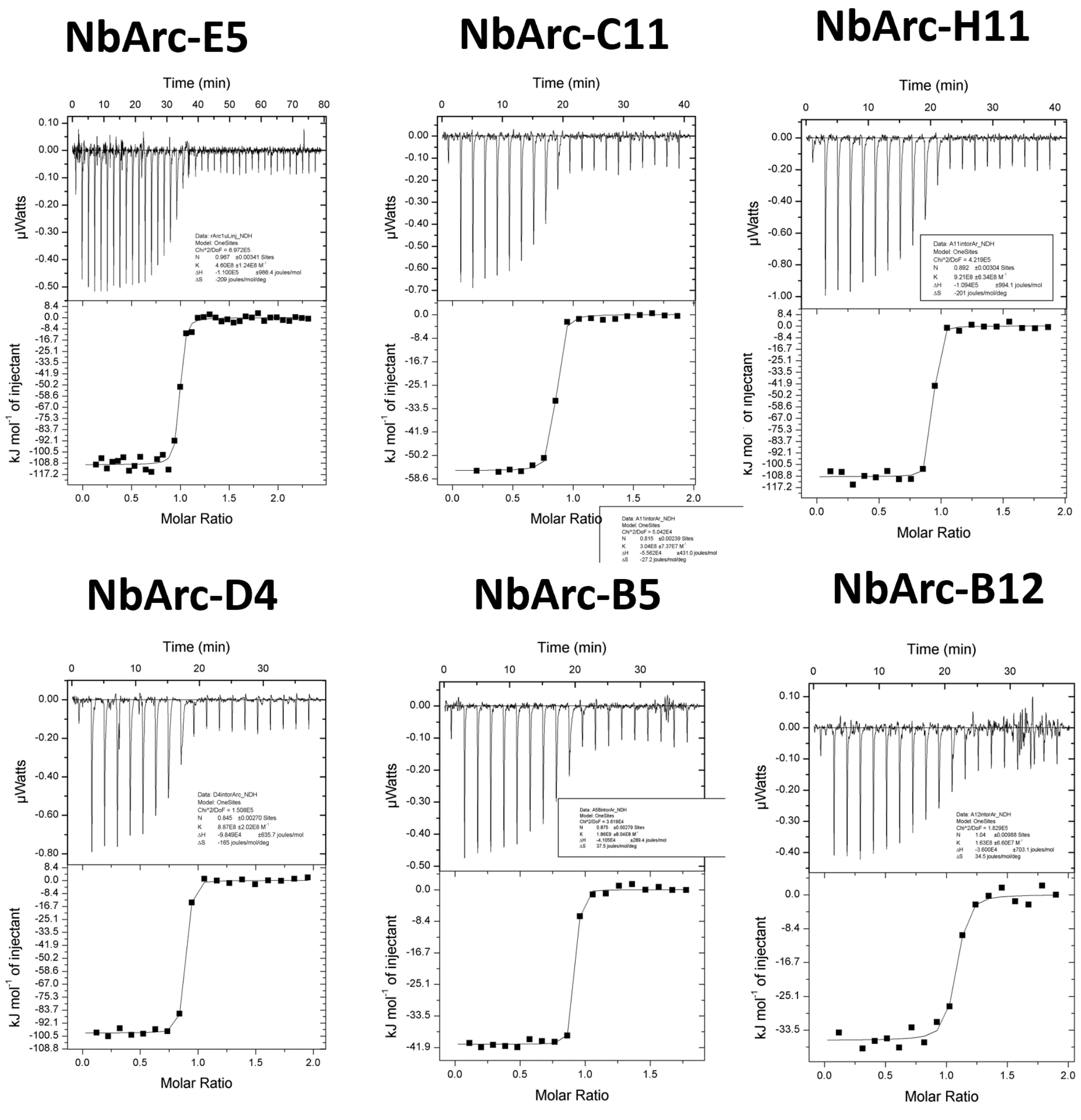


**Figure S2:** NbArc ITC titrations into FLrArc-7A, raw and integrated thermograms.

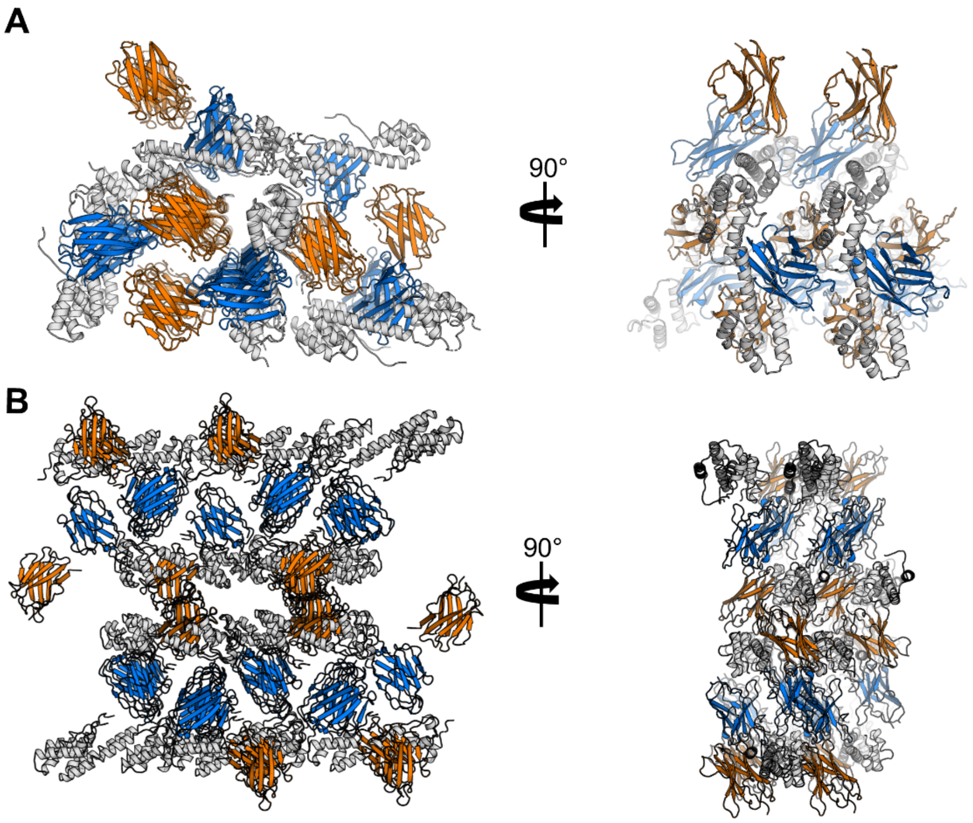


**Figure S3:** Crystal contacts in both the rArc-CTD (**A**) and the hArc-CTD extended crystal structure (**B**) were formed almost exclusively by the bound Nbs. The CTD is coloured grey, H11 orange and C11 blue.

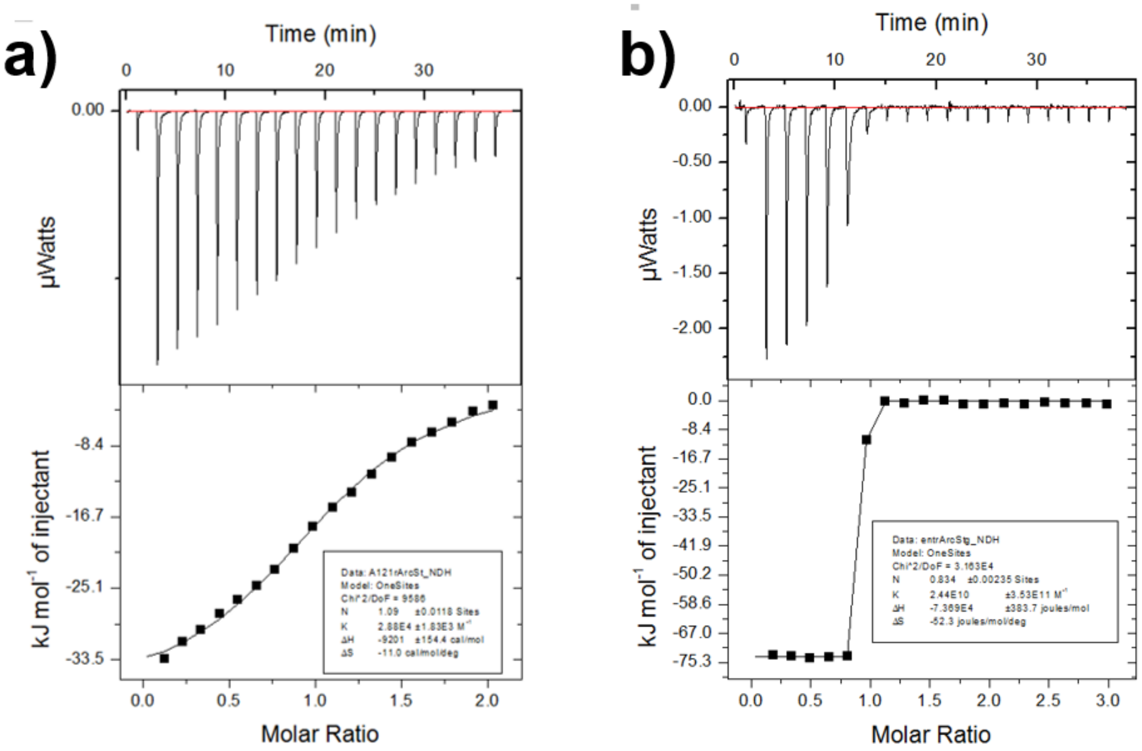


**Figure S4:** Raw ITC thermograms. **a)** Titration of the Stg peptide into FLrArc-7A and **b)** titration of NbArc-H11 into Stg-bound FLrArc-7A.

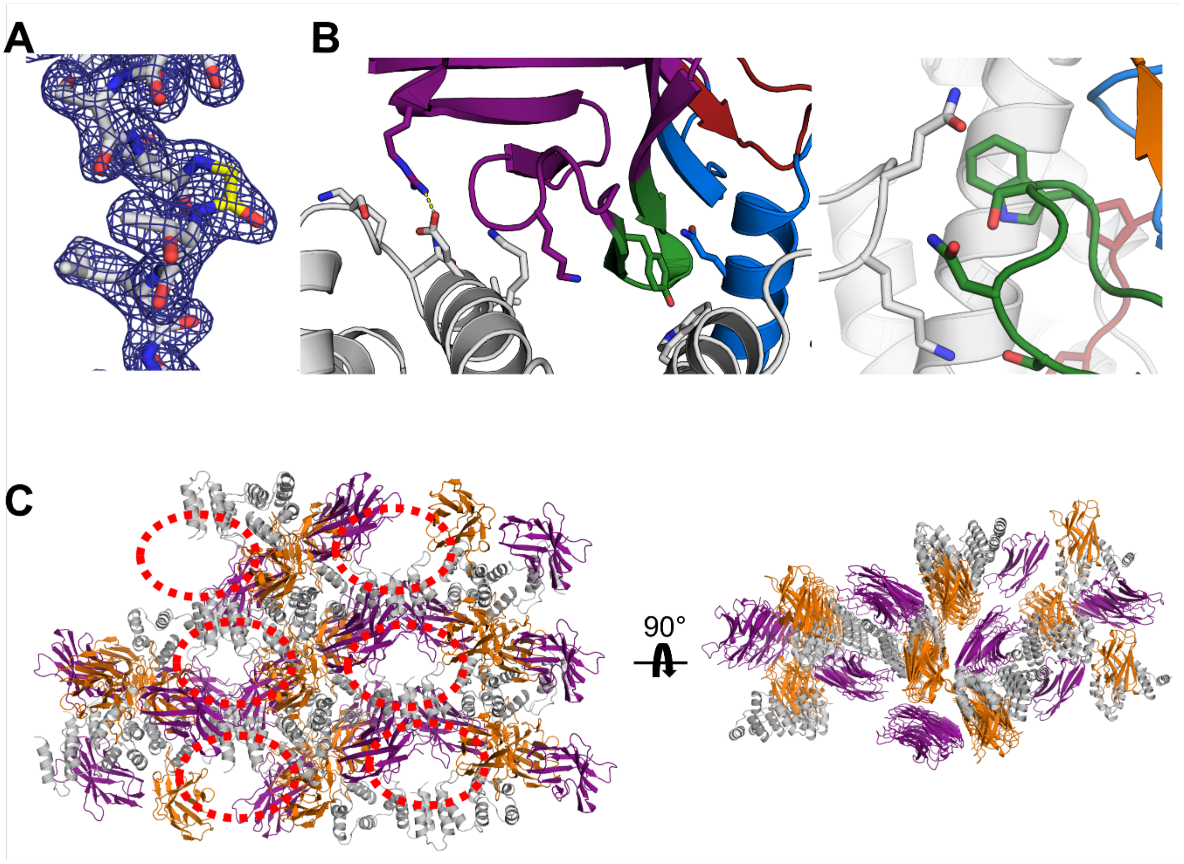


**Figure S4:** Crystal structure of the broken hArc-CTD conformer in complex with NbArc-H11 and -C11. **A** Electron density map ($2F_O$-$F_C$ at 1.5$\sigma$ in blue) of the hinge region where Gly277 is highlighted in yellow, demonstrating the lack of ambiguity in building the broken hinge region. **B** The additional Nb interactions observed in the crystal. FR region of C11 (left) binds the N-lobe and H11 (right) shows modest interactions with the collapsed C-lobe. **C** Packing in the crystal, where crystal contacts are exclusively formed by bound Nbs. Empty volumes in the lattice which might contain the unstructured C-terminal portion of the CTD are highlighted with red broken circles.

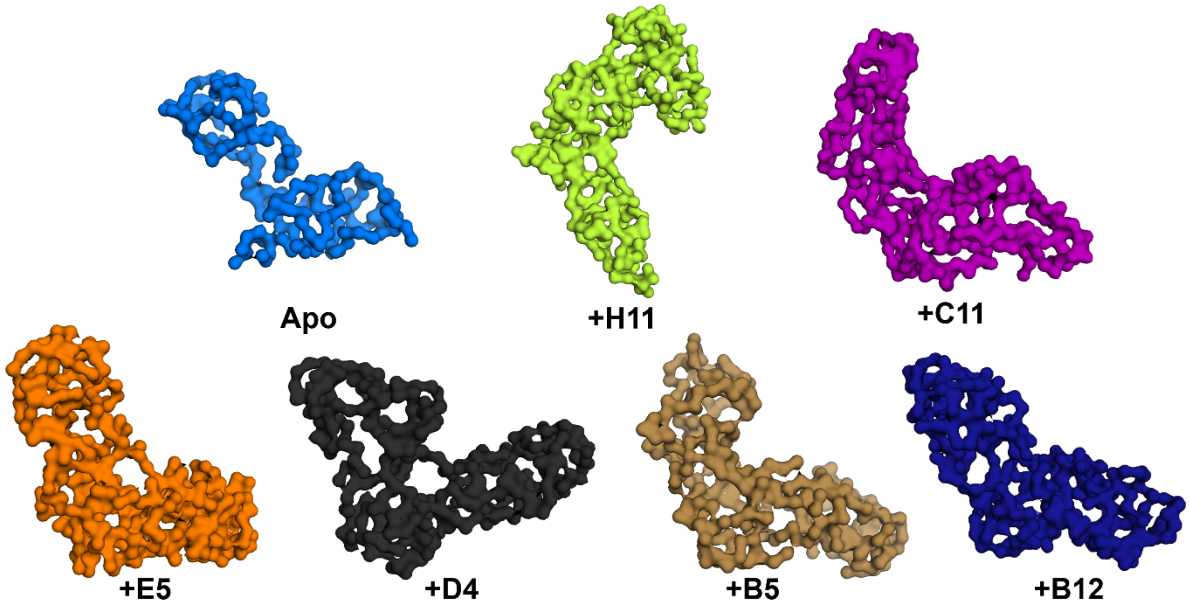


**Figure S5**: *Ab initio* models of the hArc-CTD in apo state and in complex with the anti-Arc Nbs derived from the SAXS data shown in Figure 13.

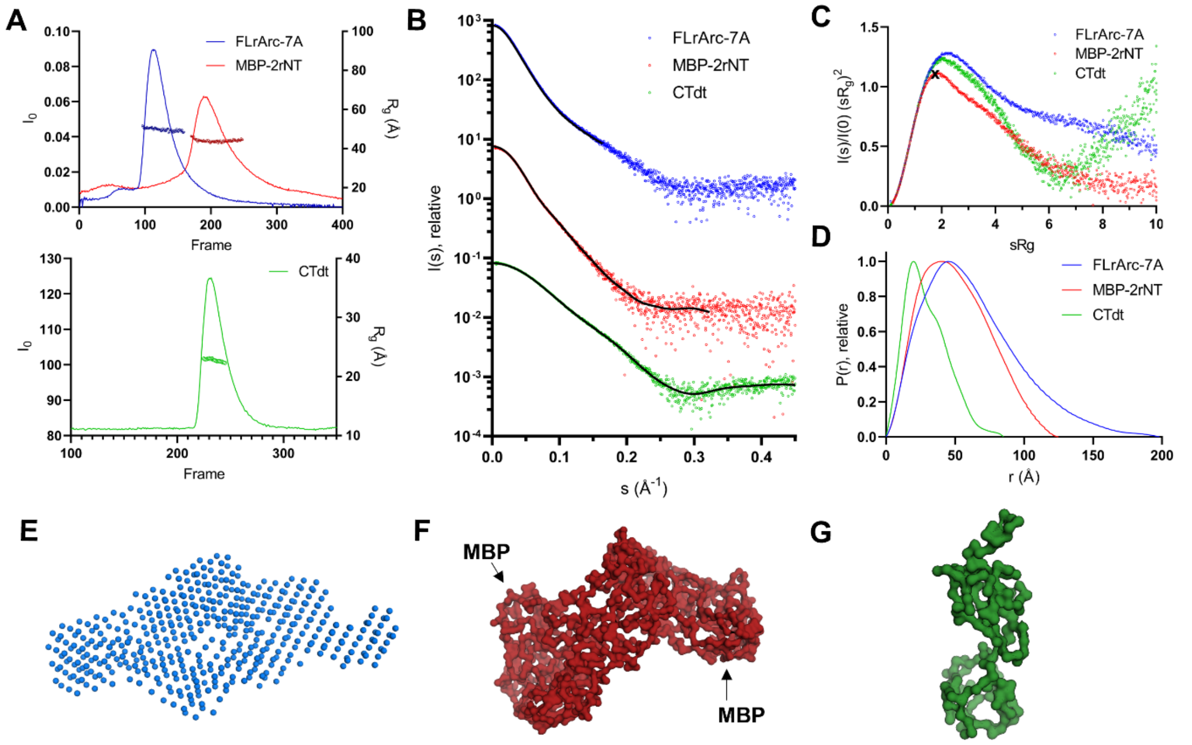


**Figure S6:** SEC-SAXS analysis of MBP-2rNT, and the other two Arc constructs used in this study. CTdt refers to the hArc-CTD construct used. **A** SEC elution profiles for the three constructs. Extrapolated I0 of each frame (lines) are plotted on the left Y-axis and open rings indicate the calculated $R_g$ of the main peak data frames used for data processing and modelling. **B** Scattering profiles of each sample. The data fits from GNOM are shown as black lines and the curves are displaced by two logarithmic units, for clarity. **C** Scattering data shown on a dimensionless Kratky plot. The X indicates the expected maximum for a fully rigid spherical particle (√3, 1.104). The rigidity of the MBP-2rNT construct was apparent. **D** Distance distribution profiles. FLrArc-7A showed a characteristic wide profile, indicating elongation, hArc-CTD showed a two-peak profile, typical of bilobar structures and the bell-shaped distribution of MBP-2rNT indicated a more compact fold. d) Ab initio model of the dimeric rArc mutant, produced with no forced symmetry in DAMMIN (χ^2=0.9992). **E** and **F** Ab initio models of the MBP-2rNT dimer and g) hArc-CTD, produced using GASBOR without forced symmetry (χ^2=1.790 for MBP-2rNT and χ^2=1.261 for hArc-CTD).
